# An aptamer-based magnetic flow cytometer using matched filtering

**DOI:** 10.1101/2020.04.14.041210

**Authors:** Chih-Cheng Huang, Partha Ray, Matthew Chan, Xiahan Zhou, Drew A. Hall

## Abstract

Facing unprecedented population-ageing, the management of noncommunicable diseases (NCDs) urgently needs a point-of-care (PoC) testing infrastructure. Magnetic flow cytometers are one such solution for rapid cancer cellular detection in a PoC setting. In this work, we report a giant magnetoresistive spin-valve (GMR SV) biosensor array with a multi-stripe sensor geometry and matched filtering to improve detection accuracy without compromising throughput. The carefully designed sensor geometry generates a characteristic signature when cells labeled with magnetic nanoparticles (MNPs) pass by thus enabling multi-parametric measurement like optical flow cytometers (FCMs). Enumeration and multi-parametric information were successfully measured across two decades of throughput. 10-µm polymer microspheres were used as a biomimetic model where MNPs and MNP-decorated polymer conjugates were flown over the GMR SV sensor array and detected with a signal-to-noise ratio (SNR) as low as 2.5 dB due to the processing gain afforded by the matched filtering. The performance was compared against optical observation, exhibiting a 92% detection efficiency. The system achieved a 95% counting accuracy for biomimetic models and 98% for aptamer-based pancreatic cancer cell detection. This system demonstrates the ability to perform reliable PoC diagnostics towards the benefit for NCD control plans.

## 1. Introduction

Noncommunicable diseases (NCDs), generally known as chronic diseases, have become the primary risk of death with unprecedented population-ageing (United Nations et al., 2017). According to the World Health Organization (WHO), more than 80% of deaths are caused by NCDs in countries where at least 20% of the population is over 60 years old (World Health Organization, 2018). Inevitably, the growing impact of NCDs necessitates changes to the contemporary healthcare system that was designed many decades ago. While the successful transformation requires management of common NCDs (*e.g.*, cancers, cardiovascular diseases, diabetes, etc.), it is predicated on effective diagnosis, screening, monitoring, and treatment. The upcoming digital-health era shifts healthcare from a sick-care response to a proactive system with personalized medicine that addresses many unmet needs of NCD management (Bhavnani et al., 2016; Li et al., 2017; Wang et al., 2018; Wannenburg and Malekian, 2015).

One of the key factors to enable the unbridled ubiquity of digital health is to transition diagnostics from centralized laboratories closer to the patient in point-of-care (PoC) settings. PoC settings integrated with health information technology, telemedicine, and portable testing provide high effectiveness, low cost, easy access, and fast turnaround. Many technologies have improved the feasibility of PoC testing using optical biosensors (Kwon et al., 2016; Zhu et al., 2013), FET-based biosensors (Afsahi et al., 2018; Wang et al., 2014), electrochemical biosensors (Aronoff-Spencer et al., 2016; Jiang et al., 2017; Sun and Hall, 2019), and magnetic biosensors (Zhou et al., 2019a, 2019b), amongst many others. PoC diagnostics offers timely detection and treatment monitoring of cancers where early detection has a huge impact on the treatment outcome and survival rate while simultaneously reducing the economic burden. Optical-based instrumentation is still the workhorse in clinical diagnostics with techniques such as flow cytometry. A flow cytometer (FCM) is an essential tool in hematology for quantitative analysis of cells with applications including identifying prognostic indicators for cancer, HIV, and other time-dependent biomarkers of disease activity (Jennings and Foon, 1997; Perfetto et al., 2004; Malcovati et al., 2007). A cellular collection system can be added to realize fluorescence-activated cell sorting (FACS), which enable high throughput and multi-parameter, quantitative cellular analysis and sorting (Julius et al., 1973). However, such instrumentation requires complex optics, lasers, and photodetectors making it hard to translate to the PoC. CMOS-based optical biosensors have integrated the necessary components into portable formats (Vashist et al., 2015; Zhu et al., 2013, 2011); however, the devices still fall short of the PoC promise due to the need for extensive sample pretreatment and bulky optics that require alignment and/or calibration.

To address cancerous cell detection for NCD control, a magnetic flow cytometer (MFC) provides an alternative platform for the multi-parametric quantification of cellular information while molding PoC-friendly settings like rapid turnaround time and miniaturization without the loss of sensitivity (Helou et al., 2013; C. Huang et al., 2017; C.-C. Huang et al., 2017; Issadore et al., 2012; Lin et al., 2016; Loureiro et al., 2011; Murali et al., 2016; Reisbeck et al., 2018, 2016; Tang et al., 2019, 2019; Zhou et al., 2017). Magnetic biosensors replace the fluorescent (or colorimetric) label with a magnetic tag. Magnetic detection has less background noise than optical measurements where issues such as photobleaching and auto-fluorescence are always present requiring sample pretreatment to remedy (Cossarizza et al., 2017; Salvati et al., 2018; Williams et al., 2017). Thus, magnetic sensing simplifies the assay procedure by greatly easing the necessary sample preparation (E. Fernandes et al., 2014; Freitas et al., 2012; Gaster et al., 2009; Osterfeld et al., 2008; Rizzi et al., 2017; Wang et al., 2015).

In this work, we developed a giant magnetoresistive spin-valve (GMR SV)-based MFC using matched filtering and a multi-stripe sensor geometry to improve detection accuracy (**Fig. 1**). At a high level, the system can be explained as follows: when an analyte (*e.g.*, a cancer cell) labeled with magnetic nanoparticles (MNPs) flows over the sensor, a change in resistance of the underlying sensor is induced. The carefully designed sensor layout creates a characteristic signature from MNPs, as shown in **Fig. 1B**, thus enabling multi-parametric measurement like optical FCMs. An array of sensors spaced along the fluidic channel extract the time-of-flight (ToF), which can be used as a proxy for the size and hydrodynamic volume of the cell. To study this effect, we have done a full force vector analysis and studied the effect of different MNP sizes and flow cell dimensions. Enumeration and multi-parametric information were successfully measured across two decades of throughput. Biomimetic constructs consisting of 10-µm polymer microspheres were used as a model system where MNPs and MNP-decorated polymer conjugates were flown over the GMR SV sensor array and detected with a signal-to-noise ratio (SNR) as low as 2.5 dB due to the processing gain afforded by the matched filtering. The matched-filtering results were compared with optical observation, showing correlation rates up to 92%. The GMR SV-based MFC achieves 95% counting accuracy in the biomimetic sample (MNP-decorated polymer spheres) and expands to aptamer-based cancer cell detection with 98% correlation to an optical FCM demonstrating the ability to perform reliable PoC diagnostics towards the benefit of NCD control plans.

**Figure 1.**
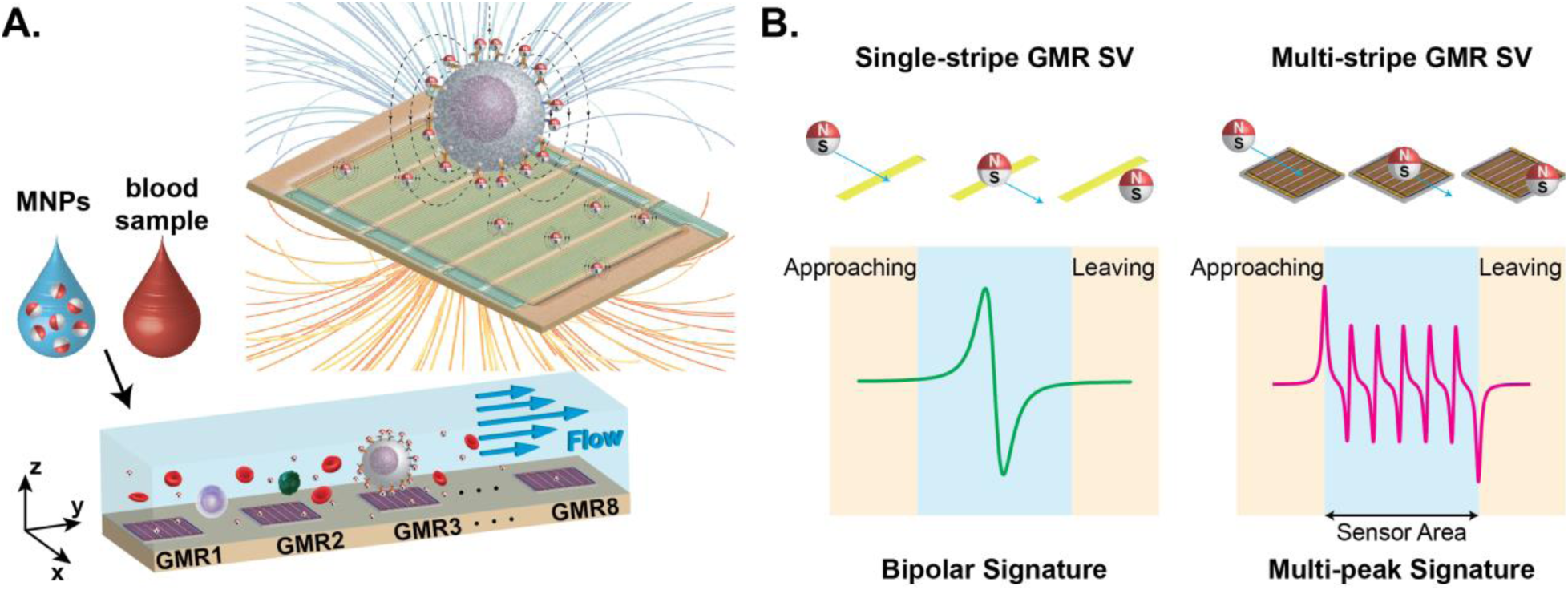
Illustration showing: **A.** Operation of a GMR SV-based magnetic flow cytometer (MFC) where MNP-decorated cells flow over GMR SV sensors. **B.** Signature from conventional single-stripe sensors with a simple bipolar-peak which increases the false detection events and the proposed multi-stripe configuration that enhances the signal differentiation by creating a unique magnetic signature.

## 2. Materials and Methods

### 2.1 Magnetic flow cytometer

The magnetic flow cytometer (MFC) consisted of a GMR SV chip (MagArray), a NdFeB permanent magnet, a microfluidic channel, and electrical readout circuitry (**Supplemental Fig. 1**). Each GMR SV chip has 80 individually addressable sensors arranged in an 8×10 matrix where each sensor is 120×120 µm^2^ on a 280 µm pitch with a nominal resistance (*R*_0_) of 1464 Ω and a mean magnetoresistance (MR) ratio of 7.99% (**Supplemental Fig. 2**). Only one row of sensors (*n* = 8) was used in this work. The NdFeB permanent magnets (K&J Magnetics, B881, B882, B882-N52, BCC2, or BCC2-N52) were mounted horizontally 4.5 mm below the sensor chip with an out-of-plane field that ranged from 0.06 to 0.13 T, as measured by a gaussmeter (Lake Shore Cryotronics, 475DSP).

The GMR SV sensors were read out using lock-in detection excited by a 1 V_pp_ sinusoidal source at 7 kHz generated by a data acquisition card (National Instruments, PCIe-6361), as shown in **Supplemental Fig. 3**. The resulting current was amplified by a transimpedance amplifier (TIA) implemented using an OpAmp (Analog Devices, AD8655) with resistive feedback (*R*_F_ = 42.2 kΩ). A bleed resistor (*R*_B_ = 1.5 kΩ) was used to cancel the non-magnetoresistive portion of the current and avoid saturating the TIA, thus enabling the gain to be increased by 28 dB. Eight parallel channels of this circuit were assembled on a custom printed circuit board (PCB). The TIA outputs were sampled at 125 kSps/ch. and processed in LabVIEW using a Fast Fourier Transform (FFT) to demodulate the signal (125-point FFT, 1 ms acquisition time). The input-referred noise of the system was measured to be 4.2 mΩ_RMS_ and spectrally white (**Supplemental Fig. 4**).

Optical measurements were taken by an optical microscope (Motic, #BA310MET-T) with a microscope camera (Moticam, #1080) or a lens-mounted mobile phone (iPhone X) for video recording under different flow rates. The videos were post processed with a monochromatic filter and magnetic analytes were enumerated using a custom written MATLAB code via size thresholding.

### 2.2 Microfluidics

Microfluidic channels were fabricated using a standard poly(dimethylsiloxane) (PDMS) process with SU-8 molding and PDMS curing with channel widths ranging from 90 to 120 μm and heights ranging from 14 to 40 μm. GMR SV chips were placed in a UV-ozone chamber (UVOTECH, HELIOS-500) for 15 minutes prior to bonding with the PDMS microfluidic channels. The microfluidic chips were subsequently aligned and cured for 1 hour at 75 °C. The inlet and outlet of the PDMS channel were mechanically drilled and connected to a syringe pump (NE-300, New Era Pump Systems) with poly(tetrafluoroethylene) (PTFE) tubing.

### 2.3 Magnetic nanoparticles

Superparamagnetic MNPs, Dynabeads M-450 (Invitrogen, #14011), Dynabeads M-280 (Invitrogen, #11205D), Bio-Adembeads (Ademtech, #03121), and SHS-30 (Ocean NanoTech, #SHS-30-01), were used in all experiments with hydrodynamic diameters of 4.5 μm, 2.8 μm, 200 nm, and 40 nm, respectively. Dynabeads M-450 and Dynabeads M-280 with a core particle size of 7.7 nm were washed 3× with 0.1× PBS before diluting to 1:400 and 1:650, respectively. A nonionic surfactant, Tween 20 (Sigma-Aldrich, #P1379), was added to the diluted Dynabeads solution at a dilution of 0.05%. Streptavidin-coated Bio-Adembeads and SHS-30 were centrifuged each time before the washing step (the same procedure as the Dynabeads); the final dilution ratios were 1:20 and 1:1, respectively.

### 2.4 Hydrodynamic analysis

Hydrodynamic forces were calculated to determine the optimal channel height and magnetic field (De Palma et al., 2007; Liu et al., 2009; Wirix-Speetjens et al., 2005). Forces acting on a MNP included the drag force (***F***_**D**_), magnetic force (***F***_**M**_), gravity (***F***_**G**_), and DLVO forces (Van der Waals force (***F***_**VDW**_), and electro-repulsive force (***F***_**el**_)), and Langevin force (***F***_**lang**_), as shown in **Supplemental Fig. 5**. The drag force is calculated as

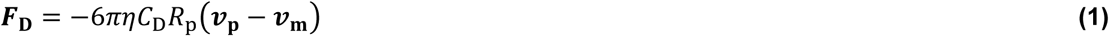

where *η* is the viscosity of the medium, *C*_D_ is the drag coefficient defined by the MNP size and shape, ***v***_**p**_ is the MNP’s velocity, and ***v***_**m**_ is the medium velocity. The magnetic force is calculated as

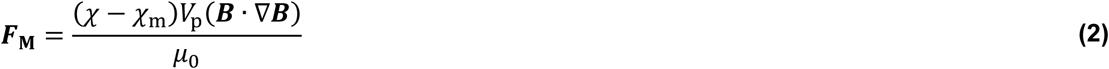

where *χ* is the volume susceptibility (dimensionless), *χ*_m_ is the medium’s volume susceptibility, *V*_p_ is the MNP volume, ***B*** is the magnetic flux density, and *μ*_0_ is the permeability of free space. Gravity is calculated as

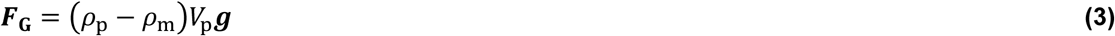

where *ρ*_p_ is the MNP density, *ρ*_m_ is the medium density, *V*_p_ is the MNP volume, and ***g*** is the gravitational acceleration. Van der Waals, electro-repulsive and Langevin forces are calculated by

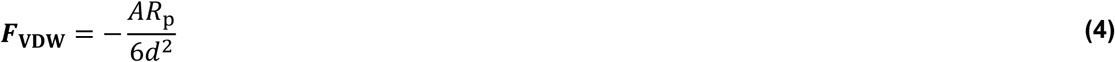

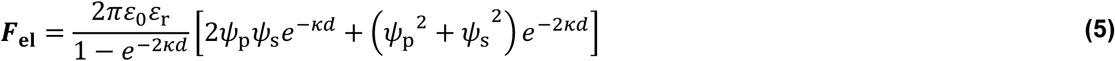

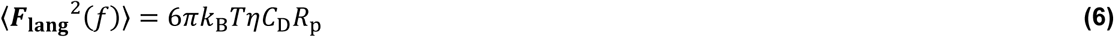

where *A* is Hamaker constant, *R*_p_ is the MNP radius, *d* is the distance between MNPs or between the MNP and sensor, *ε*_0_ is the permittivity of free space, *ε*_*r*_ is the relative permittivity, *ψ*_p_ is the MNP surface potential, *ψ*_s_ is the sensor surface potential, *κ* is the Debye–Hückel length, *k*_B_ is the Boltzmann constant, and *T* is the temperature (in Kelvin). Custom written MATLAB code was used to calculate the resulting forces based on Eqs. (1)-(6).

### 2.5 Biomimetic polymer microspheres

Biotin-coated 10-μm polymer microspheres, ProActive CP10N (Bangs Laboratories, #CP10000), were conjugated with Bio-Adembeads to create a biomimetic construct used during algorithm development and evaluation. To build such construct, an aliquot of ProActive CP10N was washed with 10× volume of wash buffer (0.1× PBS + 0.05%Tween20, pH = 7.4) three times. The pellet in the wash buffer was resuspended to 1:20 dilution. The diluted Bio-Adembeads (1:20) were added to this solution. The magnetic conjugates were formed and incubated at room temperature (18-25 °C) for 30 minutes with gentle mixing. The sample was resuspended in 20× volume of wash buffer prior to injecting into the microfluidic channel using a syringe pump (New Era Pump Systems, NE-1000).

### 2.6 Micromagnetic modelling

The Stoner–Wohlfarth model was used for magnetic modelling (Guanxiong Li and Wang, 2003). MNPs are assumed to be Langevin spheres in the field, have a linear superparamagnetic response, and give rise to dipole fields. Here, we considered only the spatially averaged magnetic field emanated from a single MNP being magnetized by the applied field (***H***_**A**_). Thus, the average field that acts on the free layer of the GMR SV sensor 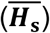 is:

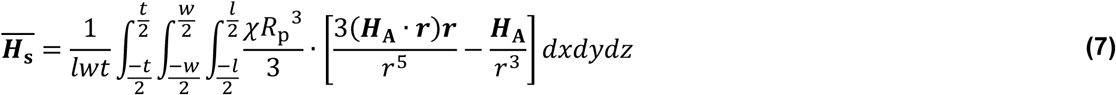

where *l* is the sensor length, *w* is the sensor width, *t* is the free layer thickness*R*_p_, ***H***_**A**_ is the applied magnetic field, ***r*** is the distance between the MNP (*x*_0_, *y*_0_, *z*_0_) and the point of free layer (*x, y, z*), ***x*** and ***y*** are the in-plane axes, and ***z*** is the out-of-plane axis, as shown in **Fig. 2A**. It is assumed that *H*_A_ points in the ***z***-direction without divergence of the in-plane component and the 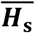 component along the long-axis (***x***, in **Fig. 1A**) is neglected due to the sensor’s insensitivity of the long-axis field. Consequently, the average field along the short-axis, ***y***, is:

**Figure 2.**
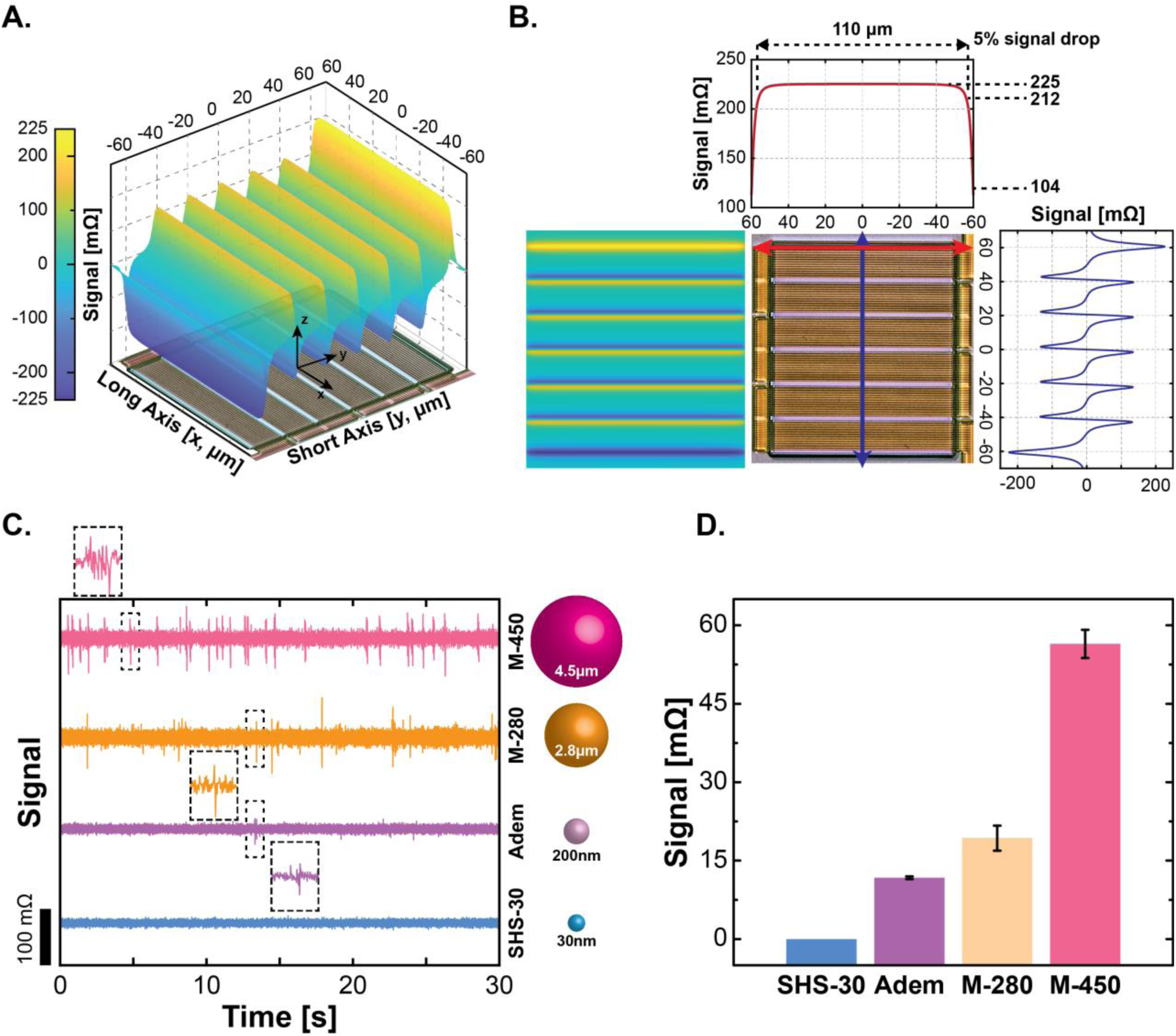
Simulation and measurement results for MNPs passing over a magnetic sensor. **A.** 3D illustration of simulated signal. **B.** 2D profile of positional signal dependence. **C.** Measurement of different MNPs and their magnetic signatures under the same flow rate. **D.** Plot of measured signal amplitude vs. MNP size.

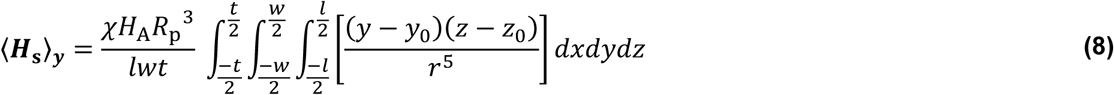

All micromagnetic simulations, such as those shown in **Fig. 2**, were performed using custom written MATLAB code implementing Eq. (8).

### 2.7 Signal processing

Cross-sensor correlation and matched filtering (MF) were applied on the acquired data to calculate the time-of-flight (ToF) across the sensor array and improve the SNR, respectively. Cross-sensor correlation involves convolving a signal segment from a detected event with the signal measured on a subsequent sensor (*i.e.* using the signature observed on Sensor 1 on Sensor 2). The resulting signal is thresholded to find the delay between the two events and the ToF is calculated based on the known sensor-sensor spacing and time difference. Matched filtering convolves the measured signal with a template. Three different templates were evaluated with matched filtering: simulation-based matched filters (SMF), energy-detection matched filters (EDMF), and previous-event matched filters (PEMF). The SMF utilizes Eq. (8) to generate a library of templates. The EDMF template quantizes the expected signature into a tertiary square waveform. Lastly, the PEMF relies on the signature detected at a previous sensor. These templates are illustrated in **Supplemental Fig. 6**. All signal processing was performed in MATLAB using custom written code.

### 2.8 Aptamer-based MFC assay

Panc-1 and MiaPaCa-2 pancreatic cancer cell lines were grown to 80% confluence in Dulbecco’s modified Eagle medium (DMEM) (Gibco, #11965084) with 10% fetal bovine serum (FBS) (Gibco, #26140079) and 1% penicillin/streptomycin (Gibco, #15070063). The adherent cells were treated with Trypsin (Gibco, #25300062) to detach them from the tissue culture flask using standard cell culture techniques. The cell viability and the size were calculated using a Vi-CELL XR Cell Viability Analyzer (Beckman Coulter). The cells were finally washed and resuspended in PBS/MgCl_2_/CaCl_2_ (Gibco, #14040133) for the assay.

The 5′-biotinylated-E07 (anti-EGFR aptamer) was generated by performing an *in vitro* transcription reaction (DuraScribe T7 Transcription Kit, Lucigen, #DS010925), as described previously (Ray et al., 2012). 5′-Biotin-G-Monophosphate (TriLink, #N-6003) at a 20 mM (final concentration) was also added to the reaction mixture for the incorporation of 5′-biotin to the E07 aptamer (Bompiani et al., 2012). The transcribed 5′-biotinylated E07 aptamer was purified by denaturing poly acrylamide gel electrophoresis (PAGE). A biotinylated anti-EGFR antibody (R&D systems, #FAB9577B-100) was also used for comparison.

The cells (∼3×10^5^) were incubated with the 5′-biotinylated E07 aptamer at a final concentration of 100 nM in 100 µL PBS/MgCl_2_/CaCl_2_ buffer at room temperature for 30 minutes with gentle mixing. The biotinylated anti-EGFR antibody was used at a 1:20 dilution ratio under similar reaction condition. Streptavidin-coated Bio-Adembeads were used in all the cellular detection assays. The beads were centrifuged/washed 3 times in 1× PBS/MgCl_2_/CaCl_2_ and diluted with DI water to 1:10 ratio. 40 µL of the diluted Adembeads were added to 90 µL of anti-EGFR aptamer or antibody-bound cells and incubated at room temperature for an additional 30 minutes with gentle mixing. The samples with ∼1.68×10^9^ Adembeads and ∼2.4×10^5^ cancer cells were finally resuspended in 1 mL buffer prior to injecting into the microfluidics.

### 2.9 Western blot

Panc-1 and MiaPaCa-2 cells were homogenized in a radioimmunoprecipitation assay (RIPA) buffer (Thermo Fisher Scientific, #89901) containing protease and phosphatase inhibitors (Thermo Fisher Scientific, #A32959). The total protein concentration was estimated and 30 µL (10 ng/µL) of the of the cell lysate were loaded and separated by SDS–PAGE before transfer to a nitrocellulose membrane (BioRad, #1620177). Membranes were incubated with primary mouse anti-EGFR antibody (BioLegend, #933901) at 1:1000 dilution and a secondary antibody, horseradish peroxidase-conjugated goat anti-mouse IgG at 1:1000 dilution (Thermo Fisher Scientific, #31430). As a protein loading control, a primary rabbit anti-GAPDH antibody at 1:1000 dilution (Abcam, #ab181602) and a secondary antibody, horseradish peroxidase-conjugated goat anti-rabbit IgG 1:1000 dilution (Thermo Fisher Scientific, #31460) was used. The antigen-antibody complexes were detected by the ECL system (Thermo Fisher Scientific, #32106). A pre-stained molecular weight marker was run in parallel to determine the molecular weight of the proteins (BioRad, #1610375).

### 2.10 Optical flow cytometry

Panc-1 and MiaPaCa-2 cells were grown to 80% confluence and treated with Trypsin to detach them from the tissue culture flask. The cell number and viability were counted as described previously. A streptavidin-phycoerythrin (SA-PE, Prozyme) fluorophore was used to label the biotinylated anti-EGFR aptamer and the antibody. Cells (∼ 1×10^6^) were first incubated with the fluorophore labeled aptamer (100 nM final concentration) or the antibody (1:20 dilution) in 100 µL PBS/MgCl_2_/CaCl_2_ for 30 minutes at room temperature. The stained cells were subsequently washed 3 times with 200 µL of PBS/MgCl_2_/CaCl_2_, resuspended in 500 µL of the same buffer and analyzed using FACSCalibur (BD Biosciences). The flow cytometry data was analyzed by using FlowJo software (BD Biosciences).

## 3. Results and Discussion

Several limitations today restrict the portability of FCMs. First, conventional optical-based FCMs require extensive sample preparation, such as cell lysis and/or matrix purification to properly detect cells/cell surface receptors (*e.g.*, CD4, EGFR, EpCAM) from crude samples due to the substantial optical background that the matrix presents (Issadore et al., 2012; Reisbeck et al., 2016). Second, FCMs often use sheath fluid to center the analytes in the middle of the channel with laminar flow and hydrodynamic focusing. Lastly, the readout instrumentation requires complex optics, lasers, and photodetectors making it hard to directly translate to PoC settings. To enable PoC, sample-to-answer operation, we minimized the amount of sample preparation required without significantly affecting the throughput or sensitivity. We accomplished this objective using two techniques: 1) switching from an optical-based to a magnetic-based readout, and 2) co-optimizing the size of the sensor to remove the need for sheath fluid while generating a complex signature that enables advanced signal processing techniques to improve the SNR.

### 3.1 Hydrodynamic focusing

The background signal in a MFC is near zero as biological samples intrinsically lack magnetic material (Gaster et al., 2009; Osterfeld et al., 2008) thus removing the need for purification steps. Rather than using magnetic guides to focus the cells over a small sensor (Helou et al., 2013; Loureiro et al., 2009b; Reisbeck et al., 2016), we use the microfluidic channel to confine the cells over a much larger sensor. Using a large sensor negatively impacts the sensitivity, but, more importantly, ensures that the flowing analytes always travel across the active sensing area removing the need for sheath flow and minimizing false negative events. As will be described later, the larger sensor enables a complex signature to be generated rather than a simple bipolar peak, as shown in **Fig. 1**.

To enable high throughput detection in this relatively small channel (120 µm) compared with other MFCs (Helou et al., 2013; Murali et al., 2016; Reisbeck et al., 2018, p. 22, 2016; Tang et al., 2019), the seal between the sensor and the microfluidic channel needed to be improved to increase the flow rate and subsequently pressure. We achieved good sealing by applying UV-ozone treatment prior to bonding the sensor chip with the PDMS, post-curing to improve the contact, and spring-clamping to mechanically intensify the sealing while maintaining the sensor reusability, as shown in **Supplemental Fig. 7**. We performed hydrodynamic analysis for several different sized MNPs to determine the optimal flow rate and magnetic field strength to balance the MNP in the middle of the channel height. Many forces were considered, including the drag force, magnetic force, gravity, and particle-particle (or particle-substrate) interactions through Van der Waals, electro-repulsive, and Langevin forces. After careful analysis, it was determined that drag force is the major contributor to the MNP’s movement in the microfluidic channel. A plot showing these forces as a function of MNP size is shown in **Supplemental Fig. 8**. The drag force is at least one order of magnitude larger than magnetic force for the largest MNP (M-450, 4.5 μm) with our pumping setup. When sub-micron-sized MNPs (Adembeads, 200nm; Nanomag-D, 130nm; and SHS-30, 40nm) are considered, magnetic force becomes comparable to DLVO forces and Langevin force. Drag force was kept dominant over other forces to allow the sample to flow in the middle of the channel and thus extract multi-parametric information.

### 3.2 Magnetic signature

Based on the average magnetic field exerted on a GMR SV sensor (as described in the Materials and Methods), the position-dependent signal of an M-450 MNP located at the sensor surface was simulated and is illustrated in **Figs. 2A** and **2B**. The magnetoresistive (MR) signal exhibits a strong dependence on the ***y***-position, while it is rather insensitive to the ***x***-position. The signal induced across the ***x***-axis of the sensor varies by only 5% from the center to the edge whereas the path along the ***y***-axis of the sensor generates a complex, multi-peak waveform with 2 major peaks and 5 minor peaks due to the serpentine sensor geometry consisting of 6 sensor stripes. This is in significant contrast to the simple bipolar-peak signal from a conventional single-stripe sensor (**Fig. 1B**) that can easily be mistaken for noise resulting in a false negative (missed detection event). This unique property will be exploited later to improve the SNR through signal processing techniques.

To evaluate and characterize the GMR SV-based MFC, we measured commercial MNPs with hydrodynamic sizes varying from 30 nm to 4.5 μm. The pumping rate for all MNP was 10 μL/min through a 120×25 μm^2^ channel. The smallest MNP, SHS-30, had no distinguishable signal (**Fig. 2C**), as expected from simulation, due to its small magnetic moment and the strong particle-particle repulsion that prevents aggregation. While we did see occasional signatures from the other sub-micron MNP, Bio-Adembead, these were likely aggregates – not signal from individual particles. On the other hand, the M-280 and M-450 MNPs induced many signals. The velocity of the M-280 is faster (due to using the same pumping rate) and the signature degrades into a bipolar peak while the M-450 demonstrates the complete signature due to the slower velocity. It should be noted that the M-280 does show the full characteristic signature at slower pumping rates, but this pumping rate was chosen to allow all particles to use the same magnetic field for fair comparison. **Fig. 2D** shows the average signal amplitude where an event is counted as anything larger than 5σ the noise level of a negative control (0.1× PBS) experiment. As expected, the smaller MNPs generated a smaller signal. It should be noted that while the out-of-plane magnetic field can be increased further to improve the amplitude, it is a delicate balance because as little as a 5-degree tilt between the sensor and a magnet can saturate the sensors. Furthermore, the divergence of the magnetic field modulates the amplitude across the sensor array. As a result, only the sensors located in the middle of the array were used for this comparison.

We also varied the channel height and the out-of-plane field to balance the magnetic force to study the effect of flowing height on signal. For example, a 130 mT field was used for tall channels (40-μm) while 60 mT was used for 19-μm channels. The measured data from the M-450 MNPs is in excellent agreement with simulation (**Supplemental Fig. 9**), while the M-280 data deviated from the simulated values, likely due to aggregation and/or chaining. As expected, small channels ensure the close proximity of the MNPs; however, aggregation and clogging were issues for channel heights of less than 12 μm and widths less than 50 μm. As such, we implemented the taller channel for rigid analytes where the channel height is at least 2× larger than the MNPs or biomimetic constructs, and comparable channel height for cells that have more shape flexibility.

### 3.3 Time-of-flight measurements

Raw data with no signal processing was collected while flowing 4.5 μm MNPs. As can be seen in **Fig. 3A**, each sensor in the linearly spaced array shows a time-sequenced response as the MNPs pass over the sensors. From these data, both the intra-sensor ToF (time between peaks within a signature) and the cross-sensor ToF (time between signatures on adjacent sensors) can be calculated. The ToF data directly measures the analyte velocity and serves as a proxy of its size, and the amplitude of the signature can be used to retrieve the vertical position of the analyte from the simulation library (**Supplemental Fig. 6**). At the current sampling rate, this system can handle velocities up to 7 mm/s without distorting the signature shape. Thus, the ToF data enables the MFC to have multi-parameter analysis (*e.g.*, height, velocity, # of MNPs) of the analyte to further aid in discrimination and reduce false positives by signal processing.

**Figure 3.**
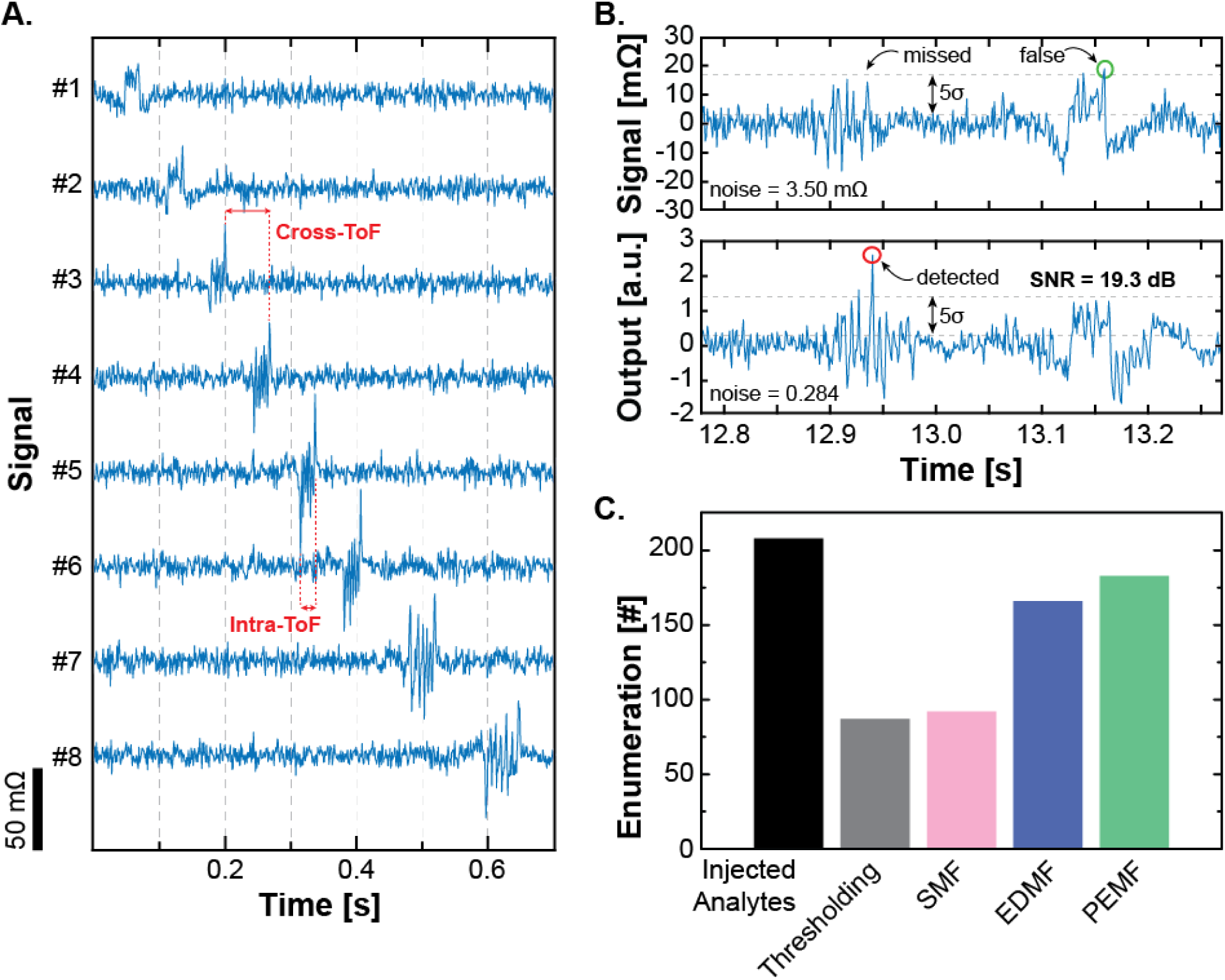
**A.** Measured data from the 1×8 sensor array exhibiting sequential signaling which enable ToF calculations and subsequent signal processing. **B.** Measured data showing missed detection using only raw-data thresholding and correct detection using matched filtering (PEMF). **C.** Comparison of enumeration using different signal processing techniques.

### 3.4 Matched filtering

Matched filtering (MF) was applied on the acquired data to improve the SNR and improve the detection efficiency (C. Huang et al., 2017). By using a multi-stripe GMR SV configuration that creates a more complex signal, the benefit of matched filtering becomes more significant compared to many previous designs that used only a single stripe sensor resulting in a simple bipolar signature (Loureiro et al., 2009b). The complicated multi-peak signature here provides a more reliable and robust matching sequence reducing the minimum detectable SNR from 14 to 2.5 dB. **Fig. 3B** shows a snippet of measured data where simple thresholding at 5σ (SNR = 14 dB) results in missing the event; however, when applying matched filtering, the event is clearly visible at 12.95 s. Also shown is a motion artifact caused by a large impulse response at 13.15 s that would be counted as a false detection with thresholding. However, since this impulse does not possess a signal-like signature, the matched filtered output is kept within the 5σ threshold and it is correctly rejected.

Three types of matched filters were evaluated: simulation-based matched filters (SMF), energy-detection matched filters (EDMF), and previous-event matched filters (PEMF). The SMF looks for measured data with a similar pattern to those in a pre-computed simulation library based on Eq. (2). The EDMF template quantizes the expected signature into a tertiary square waveform: a positive level, a zero level, and a negative level. The tertiary template created rectangular notches in a range where peaks were expected to be, roughly matching the waveform in a low-resolution fashion and allowing for tolerance in peak position. By arranging the levels in a pattern that corresponded to the expected pattern, the filter finds expected peaks while normalizing the expected peak values to minimize that degree of uncertainty. The PEMF uses signatures from upstream (*i.e.* a previous sensor) recorded events. After finding the signal sequence on any of the other sensors, the detected signature is used as a template to compare against all other sensors for correlation within a time window based on the flow rate. In a relatively short time window, the flow rate, MNP distance to the sensor, and other slowly changing environmental parameters can be regarded as constant. Therefore, each of the sensors produces their own signature but delayed in time based on the velocity. An event was claimed if the MF output exceeds the threshold which was set as 5σ from the noise. A majority voting algorithm with the eight sensors is used to reduce uncorrelated noise and declare a detection event.

Each of these matched filter templates has advantages and disadvantages. For example, owing to variations in particle size, even with the same pumping rate, particles can move at varying speeds, creating variations in the length of the target event waveform. The SMF struggles with intricate time warping between the measured signal and the template signal. In this case each peak start and end time could be slightly off from the template waveform, creating a complex warp from target to template where some areas of the signal are stretched and compressed at different rates. To deal with general differences in waveform size, we expanded the SMF library with a linear succession of filter lengths to discover each specific time variation. Stretching to different times required down-sampling or up-sampling. However, this increases the computation time significantly as the library expands. Alternatively, we could have used more advanced signal processing techniques such as dynamic time warping (DTW) (Vullings et al., 1998) but did not do so because of the high computational cost. The EDMF is elegant in that it is essentially just looking for the coarse pattern but does not have as significant improvement in the counting efficiency. Time variations were much easier to tolerate because waveform integrity was not as important for what is essentially a complex square signal. Since peaks were already being detected in a range of times rather than at a specific time, the malleability of the template proved to be much more useful than using a simulated template. Though EDMF has more flexibility in time warping and shape distortion than SMF, the velocity must be restricted to keep the intact or semi-intact complex signature. The PEMF is much more tolerant of dynamic changes in velocity. The enumeration efficacy using different signal processing techniques can be seen in **Fig. 3C**. The PEMF outperforms the other techniques and is used for the remainder of this work.

### 3.5 Optical microscopy correlation

To evaluate the efficacy and quantify the accuracy, we correlated the measured electrical signal with video recorded by an optical microscope while flowing biomimetic complexes (10-µm polysterene spheres decorated with Adembeads) and M-450 MNPs over the sensors. The Adembeads were chosen because they did not produce significant signal when not aggregated allowing differentiation between analytes bound with MNPs and unbound MNPs. **Fig. 4** shows the recorded electrical signal alongside the optical images at the same timestamps. The cross-ToF was 79 ms while travelling from sensor 3 to 4 and 78 ms from sensor 4 to 5. The intra-sensor ToF can also be extracted from the time span between two edge-peaks within a sensor, resulting in a velocity of 5.09 mm/s. The PEMF excluded the fast-flowing Adembeads and clusters based on their velocity and magnetic signature. The counting efficiency was compared between thresholding (5σ) and the PEMF using the optical counting as the ground truth (**Fig. 4B**). For M-450, both techniques have similar efficacy with detection rates of 80.62% and 92.25% for thresholding and the PEMF, respectively. However, for biomimetic MNP-decorated polymer spheres, thresholding only achieves a detection rate of 43.35% compared to the 88.18% with matched filtering. This is likely due to the complex sample which contains both biomimetic spheres, individual MNPs, and clusters of MNPs. The MF correctly rejects the latter two whereas thresholding cannot differentiate. The OM correlation reinforces the results shown in **Fig. 3C** that MF exhibits strong reliability when introducing a complicated matrix.

**Figure 4.**
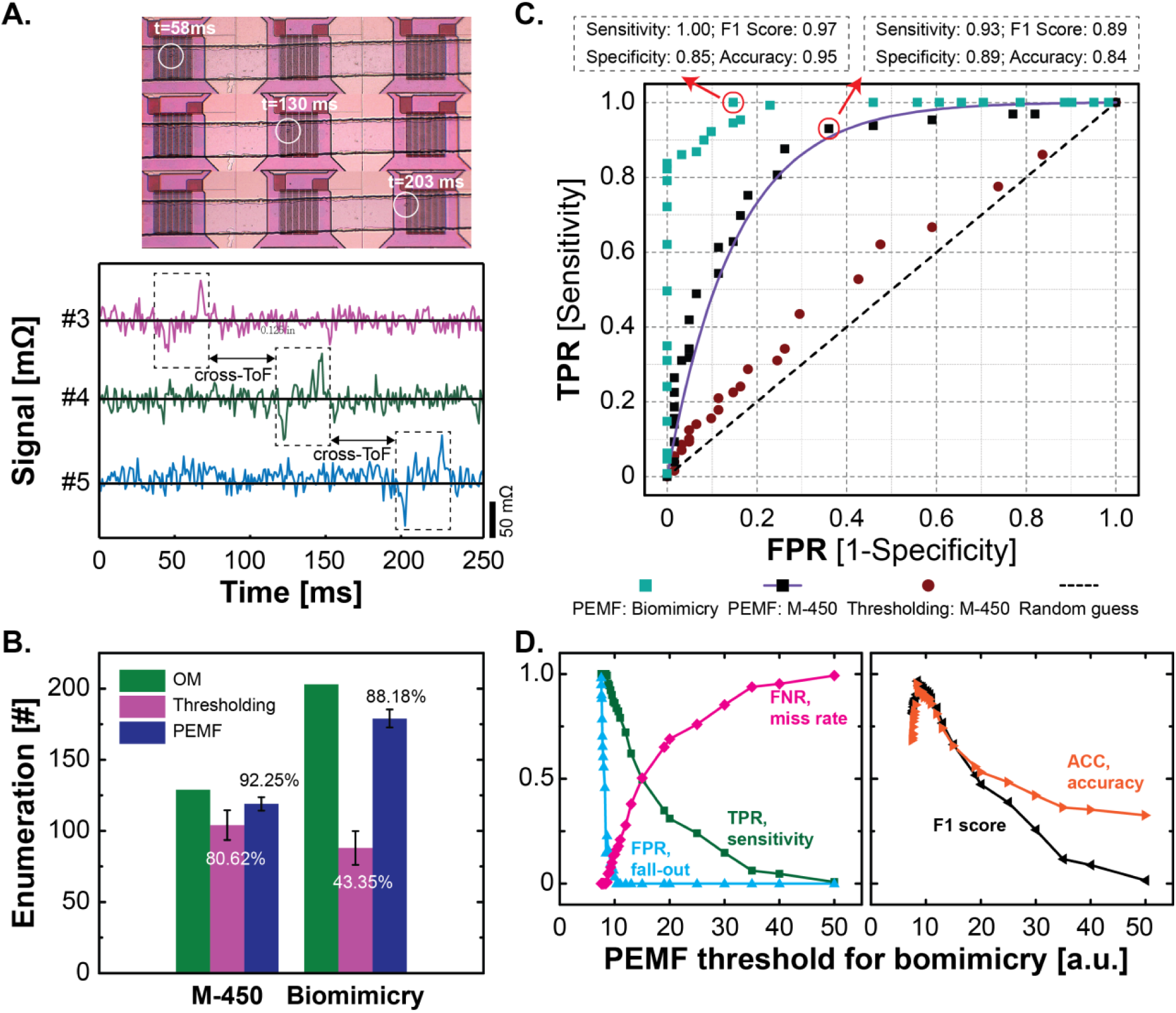
**A.** Measured real-time data of OM-monitored sensors which enable ToF measurements. **B.** Compiled event-counting data. **C.** ROC curve of the selected sensor where green dots are the biomimicry data analyzed by matched filter, square dots with the asymptote (purple curve) are the M-450 data analyzed by matched filter, red dots are analyzed by thresholding from real-time data, and the grey dashed line is the reference of random guess. **D.** Tradeoff between detection and thresholds, the highest accuracy happened when threshold was set at 8.4.

Receiver operator characteristic (ROC) curves were generated by sweeping the MF detection threshold to quantify the sensitivity and specificity of the proposed system. To establish ground truth, the flowing particles were analyzed frame-by-frame in ImageJ with size-based discrimination. Using just thresholding, the ROC curve lies very close to the random guess/chance curve, as shown in **Fig. 4C**. Applying the PEMF, the detection accuracy 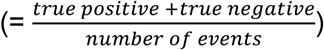 improved significantly, up to 83.68% for M-450 and 95.26% for biomimetic complexes clearly demonstrating the benefit of the MF. Different from the correlation rate in **Fig. 4B**, which was only considered the positive samples with rigid thresholds (5σ), the ROC curve reveals the tradeoff between sensitivity and specificity for a system. As shown in **Fig. 4D**, increasing the PEMF threshold improves the false negatives, while the sensitivity and false positives were reduced simultaneously. The system reached the best accuracy with a PEMF threshold of 8.4 compared to the 5σ (= 9.6) used in thresholding. This threshold strikes a balance between positive samples and negative samples but can be tuned based on the application.

### 3.6 Aptamer-based detection of pancreatic cancer cells

To establish the utility of the MFC in a cell detection assay, we used pancreatic cancer cell lines, Panc-1 and MiaPaca-2, that overexpress epidermal growth factor receptor (EGFR) on the cell surface. The Panc-1 cells had a 19.51 µm mean diameter (**Supplemental Fig. 10**). Therefore, we used the 20-μm-microfluidics channel height that is close to the size of the Panc-1 cells. A 5′-biotinylated 2′-fluropyrimidine modified RNA aptamer (E07) that binds to EGFR with high affinity and specificity was used for the cell-labeling reaction (Ray et al., 2012). We also used, a biotinylated anti-EGFR antibody as an additional cancer cell staining reagent. Western blot and optical FCM were used to verify the surface expression of EGFR on the pancreatic cancer cell lines. Panc-1 cells expressed more EGFR compared to the MiaPaca-2 cells as detected by both the western blot and optical FCM results (**Supplemental Fig. 11-12**).

Panc-1 cells were conjugated with the biotinylated anti-EGFR aptamer and subsequently with the streptavidin-coated MNPs. This complex was then injected into the microfluidic channel and measurements were collected at 0.1 µL/mL of throughput. As a negative control, we also injected PBS buffer, the MNPs alone, Panc-1 cells, and a mixture of MNPs and Panc-1 cells (without the aptamer linkers). In **Fig. 5A** and **5B**, little to no counted events were detected in the PBS buffer (*n* = 0), the Panc-1 cells (*n* = 0), or MNPs (*n* = 38) using the PEMF. A small number of counted events (*n* = 769) were detected in the Panc-1 and MNP mixture (without the biotinylated E07 aptamer linker), likely due to nonspecific binding on the cell surface from extremely excessive MNPs. However, in the presence of biotinylated E07 aptamer linker, the counted events increased nearly tenfold up to 7,140 per 5 μL in the Panc-1-E07-MNP mixture. To assess MFC performance in future applications, we further employed varying throughputs to detect Panc-1 with biotinylated E07 aptamer in culture media (**Supplemental Fig. 13**). The data was collected at rates from 0.1 to 50 µL/min, indicating the successful enumeration across two decades of throughput.

**Figure 5.**
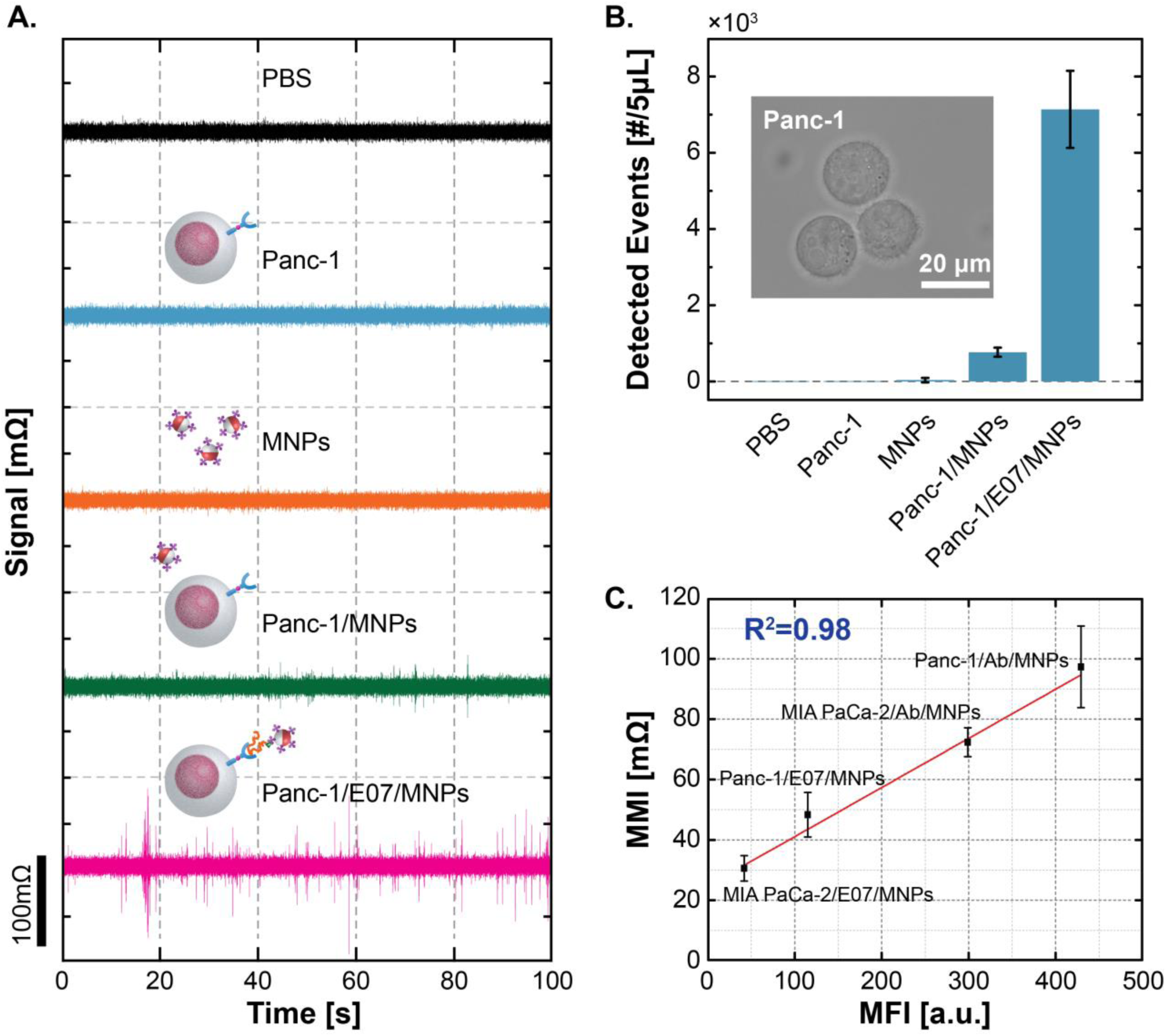
**A.** Real-time measurements of Panc-1 model study using E07 aptamer. **B.** Enumeration plot calculated from MFC measurements, and the inset shows the visualized Panc-1 cells captured by microscope. **C.** Correlation between mean magnetic intensity (MMI) and mean fluorescence intensity (MFI) with different linkers, anti-EGFR-antibody (shown as Ab) and E07 aptamer, and pancreatic cancer cell lines, Panc-1 and MIA PaCa-2. The error bars represent the counting difference throughout the measurements across 8 addressable sensors.

Lastly, we compared the MFC and optical FCM data, shown in **Fig. 5C**. Mean fluorescence intensity (MFI) was calculated from the histogram plots of the optical flow cytometry data, and plotted against the mean magnetic intensity (MMI) values that were measured from the peak amplitudes of MFC data in each detected event. A very high correlation was obtained between the two flow cytometer sensing modalities (R^2^ = 0.98). In addition, quantification of the bound MNPs can be derived from the MMI simulation (**Supplemental Fig. 14**) that indicates the amount of bound MNPs in the presence of anti-EGFR-antibody is in the range around 10^4^ per cell. Notably, the difference in the protein expression of EGFR on the cell surface of these two cell lines, Panc-1 and MiaPaca-2, were also reflected in the MFC data obtained by using two different linkers, the anti-EGFR aptamer and antibody. Taken together, the data validates our proposed method against an optical FCM that is regarded as the gold-standard instrument.

### 3.7 Comparison

There has been significant interest in MFCs over the past decade, as shown in Table 1. Most prior MFCs have used high cross-sectional area to surface area ratio microfluidics 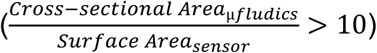 to achieve high throughput (A. C. Fernandes et al., 2014; Helou et al., 2013; Lin et al., 2014b, 2014a; Murali et al., 2016; Reisbeck et al., 2018, 2016; Vila et al., 2014). However, since the magnetic signal relies on the proximity to the sensing region (inversely proportional to the distance cubed), this is not typically a favorable design decision. As such, high area-ratio microfluidic setups lose most signals when MNP-decorated analytes flow near the middle of channel height compared to near the sensor surface. Some prior work have used magnetic chevrons (Helou et al., 2013; Reisbeck et al., 2018, 2016) or electrical current lines (Loureiro et al., 2009a) to guide the MNPs, which are close to the bottom of the channel, over the sensors and jetted successively to roll over the designated sensor area. Magnetic guides do improve the detection efficiency by focusing the analytes, however, they typically need slower flow rates which decreases the throughput and is prone to clogging. Another approach uses a strong magnetic field to attract the flowing MNP-decorated analytes near to the sensor surface, but the trajectory path makes signal modelling hard to translate into multi-parametric information and only allows binary outcomes. Furthermore, the tradeoff between forces acting in the microfluidics is complicated when using high throughput, and this kind of force analysis was mostly done for magnetic sorting (De Palma et al., 2007; Liu et al., 2009; Wirix-Speetjens et al., 2005). As such, prior MFCs have not had high detection efficiency and high throughput as needed for rare-cell detection (*e.g.*, circulating tumor cells). Considering the sensor design, miniaturization improves the sensor sensitivity, but the traditional single-stripe sensor geometry gives rise to the simple bipolar peak which is hard to differentiate signals from the noise in low SNR settings. To address this, we developed a GMR SV-based MFC with a large sensor, the size of the microfluidics, and used matched filtering to recover the sensitivity and improve the specificity. This platform uses a multi-stripe GMR SV sensor with a large active area and serpentine geometry that results in a unique signature. Our work demonstrates that the accuracy can be up to 95% in complex sample while the throughput strides across two decades that fits the clinical needs.

**Table 1.** Comparison of published magnetic flow cytometers (MFCs).

|  |  | *Lab Chip* '11 | *Sci. Transl. Med.* '12 | *Lab Chip* '13 | *SREP* '16 | *JSSC* '17 | *Biosens. Bioe.* '18 | *CICC* '19 | This work |
| --- | --- | --- | --- | --- | --- | --- | --- | --- | --- |
| Device & System | sensor | GMR | μHall | GMR | GMR | spiral transformer | GMR | LC oscillator | **GMR SV** |
| | sensor size (w*l, $\mu\text{m}^2$) | 3x40 | 8x8 | 2x30 | 2x30 | ~ 35x50 | 2x30 | 160 (diameter) | **120x120** |
|  | sensor configuration | single-stripe | cross | Wheatst one bridge | Wheatstone bridge | planar coil | Wheatstone bridge | planar coil | **multi-stripe** |
|  | sensor array | 1x3 | 2x4 | 1x4 | single sensor | 1x4 | single sensor | 7x7 | **1x8** |
| | bead size | 50 nm | 10, 12, 16 nm<br>3, 8 $\mu\text{m}$ | 200 nm<br>12 $\mu\text{m}$ | 50, 200 nm<br>4, 6, 8, 12 $\mu\text{m}$ | 1, 4.5 $\mu\text{m}$ | 6, 8, 12 $\mu\text{m}$ | 2.8, 3 $\mu\text{m}$ | **200 nm**<br>**1, 2.8, 4.5 $\mu\text{m}$** |
| Microfluidics | Dimension (w*h, $\mu\text{m}^2$) | 150x14 | 125x25 | 700x200 | 700x150 | 200x100 | 1500x150<br>1500x300 | 60000 | **90x14 – 120x40** |
|  | focusing technique | sheath | sheath | magnetic | magnetic | sheath | magnetic | -- | **laminar** |
| | throughput ($\mu\text{L}/\text{min}$) | 1.26 | 1.67 – 16.67 | 24 – 60 | 60 | 2.5 | 72 – 90 | 36 | **0.1 – 50** |
| Signal | signal processing | peak thresholding | N/A | peak thresholding | peak integral | matched filter | peak integral | cross-correlation | **matched filtering, cross-correlation** |
| Performance | OM correlation | -- | -- | Yes | Yes | -- | Yes | Yes | **Yes** |
|  | detection efficiency | N/A | N/A | N/A | N/A | 74% – 87% | N/A | N/A | **\*92.25%**<br>**\*\*88.18%** |
|  | accuracy | N/A | 0.96 | N/A | N/A | N/A | N/A | N/A | **\*83.68%**<br>**\*\*95.26%** |

\*measured with Dynabeads M-450
\*\*measured with biomimetic polymer microsphere

## 4. Conclusion

In this work, we developed a GMR SV-based MFC that innovates and evolves key areas of cellular detection related to system design and signal processing. The micromagnetic simulation and hydrodynamic force study were performed to assess the fluidics design to maximize signal. Several commercial-available MNPs were measured and compared with the 200-nm MNPs selected for biomimetic model and cellular measurements. We demonstrated the improvement in SNR when applying matched filtering and compared different template functions. The PEMF exhibited a 5.6× improvement in the minimum SNR requirement and performed significantly better than traditional thresholding in terms of enumeration. Measurement results were compared with OM-monitored enumeration achieving detection efficiencies of 92% in the bead-only assay and 88% in the biomimetic assay. ROC analysis showed the sensitivity and specificity tradeoff achieving an accuracy of up to 0.95 for the model system. To lay the foundation for rapid cancer cell detection, we used an EGFR-targeting aptamer and detected pancreatic cancer cells across two decades of throughput. The magnetic flow cytometry was compared against an optical FCM, the gold standard in cellular assay, demonstrating high correlation (98%). This work lays the foundation for the proposed PoC “*sample-to-answer*” platform needed for NCD control.

## Acknowledgements

This work was supported in part by the National Science Foundation (Grant ECCS-1454608) and Qualcomm.

## Appendix A. Supporting information

**Supplemental Figure 1.**
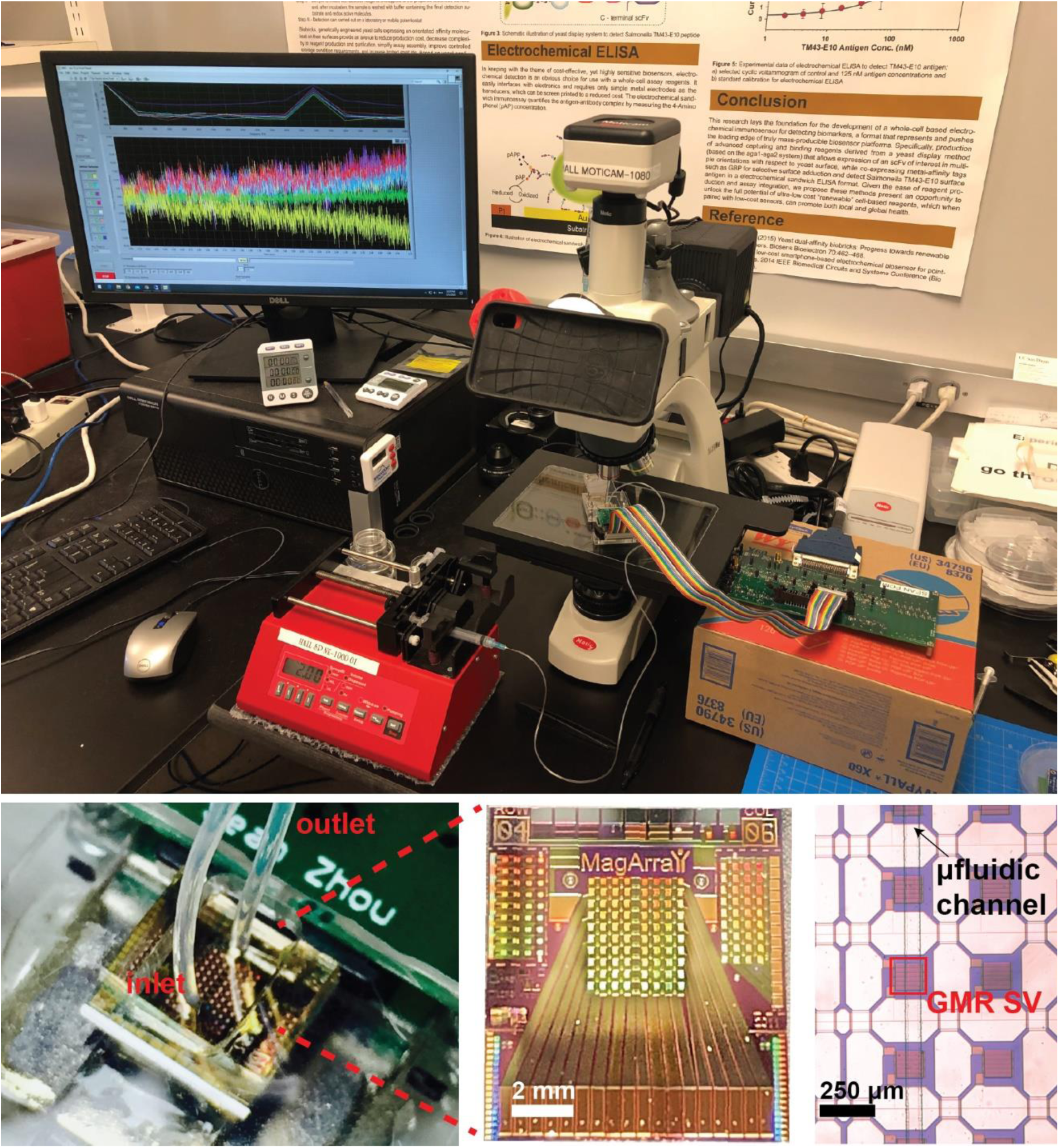
Photographs of the system (top) and zoomed-in view (from bottom left to right) of the sensor setup, a sensor chip, and the microfluidic channel. The desktop-based testing setup shows the MFC system and all components. Measurements were recorded through a custom written LabVIEW interface (as shown on PC screen).

**Supplemental Figure 2.**
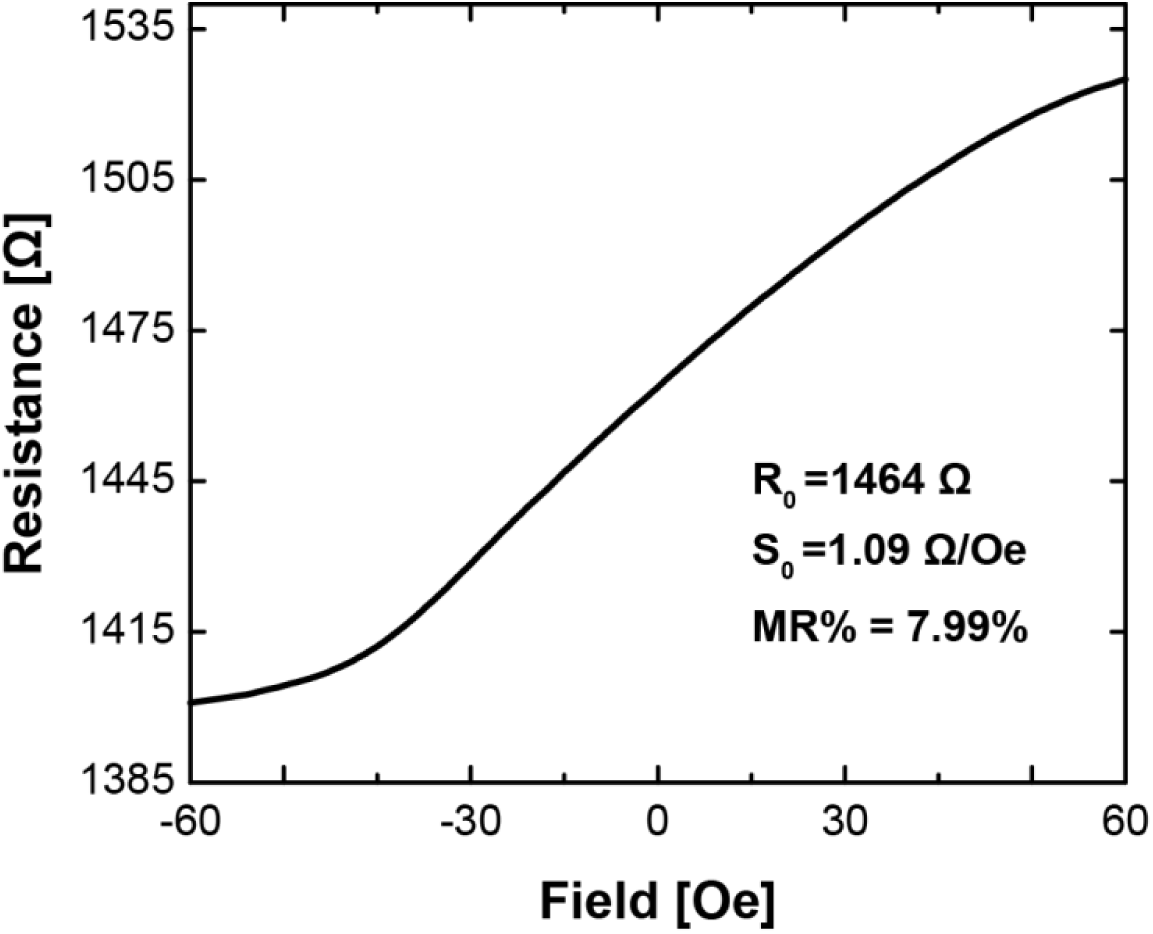
Measured magnetoresistance (MR) curve of GMR SV sensor. Each GMR SV has a nominal resistance (*R*0) of 1464 Ω, magnetoresistance (MR) ratio of 7.99%, and a sensitivity of 1.09 Ω/Oe.

**Supplemental Figure 3.**
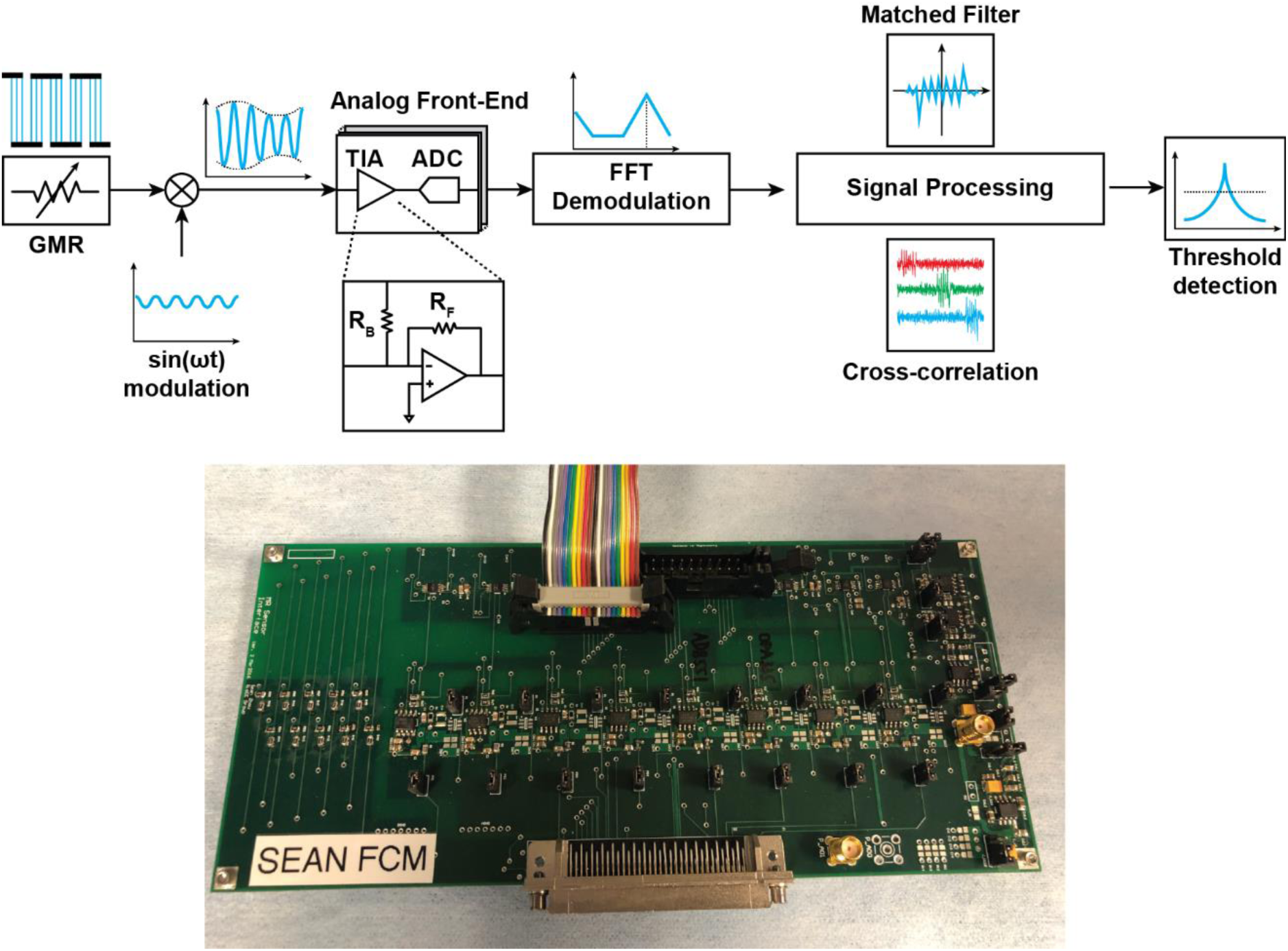
Block diagram of the measurement electronics and photograph of the PCB. The GMR sensors are modulated by a sinusoidal voltage and the resulting currents are quantized by a transimpedance amplifier (TIA) and an ADC. A bleed resistor (*R*B) removes most of the sensor baseline current. Digital signal processing performs demodulation then applies matched filtering and cross correlation to enable high SNR signal detection.

**Supplemental Figure 4.**
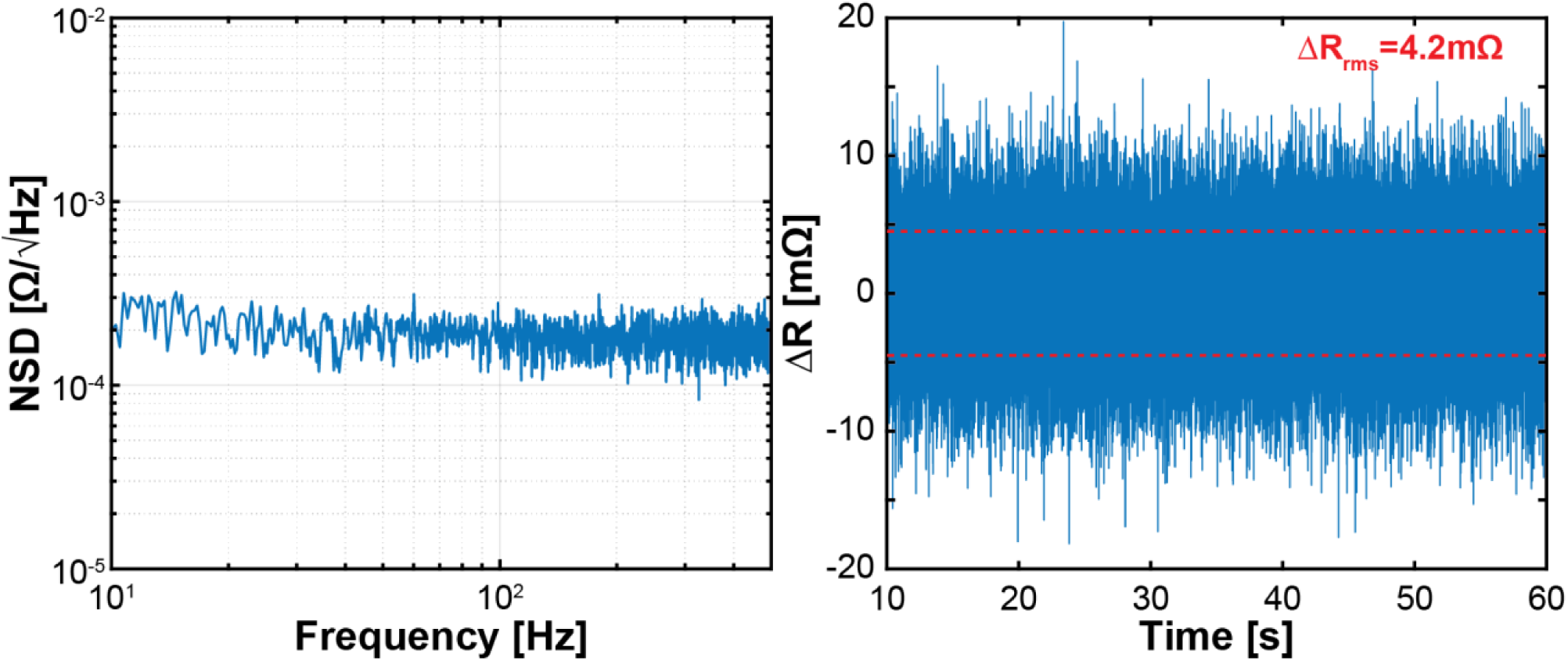
Measured spectrum and noise (left). The noise is white over the bandwidth. Measured transient noise (right).

**Supplemental Figure 5.**
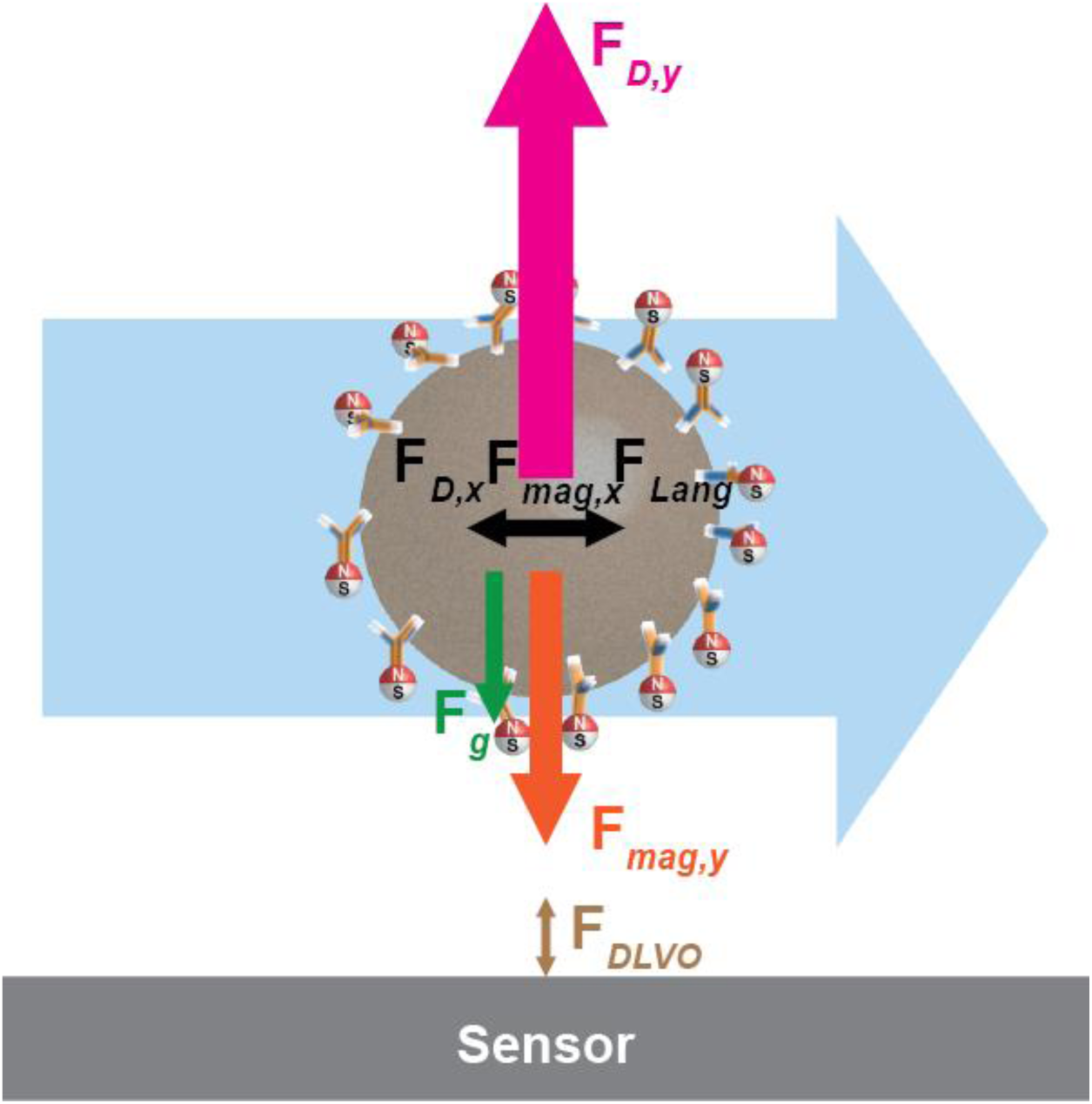
Illustration of forces acting on a magnetic nanoparticle labelled analyte. Pumping rate and corresponding flow velocity fast enough to keep drag force dominant over other forces.

**Supplemental Figure 6.**
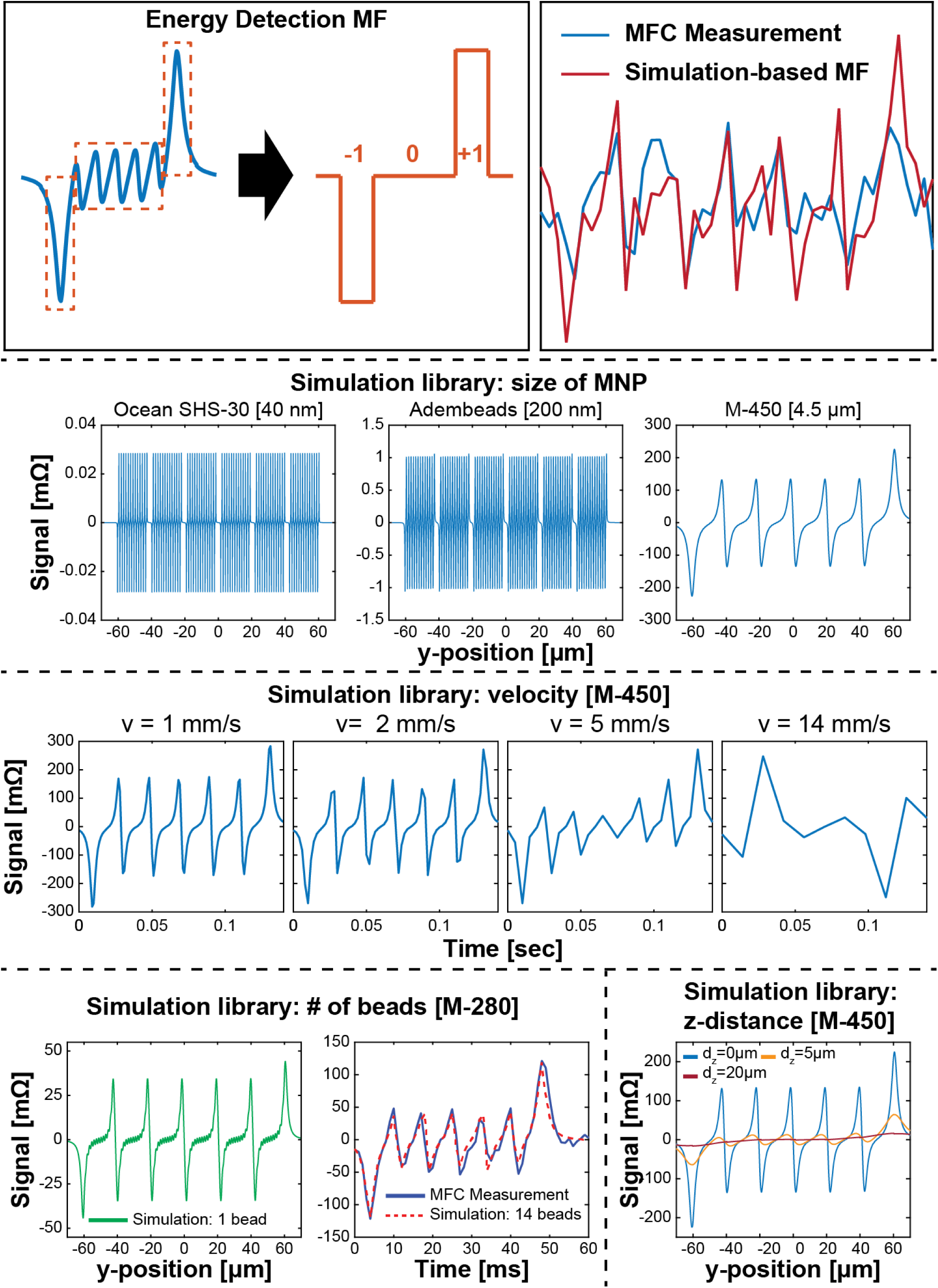
EDMF and SMF templates. EDMF digitalizes signatures into three regions: −1, 0, and 1. SMF are based on MATLAB simulations with different parameters: MNP size, flow height (distance from sensor surface), number of MNPs, and velocity.

**Supplemental Figure 7.**
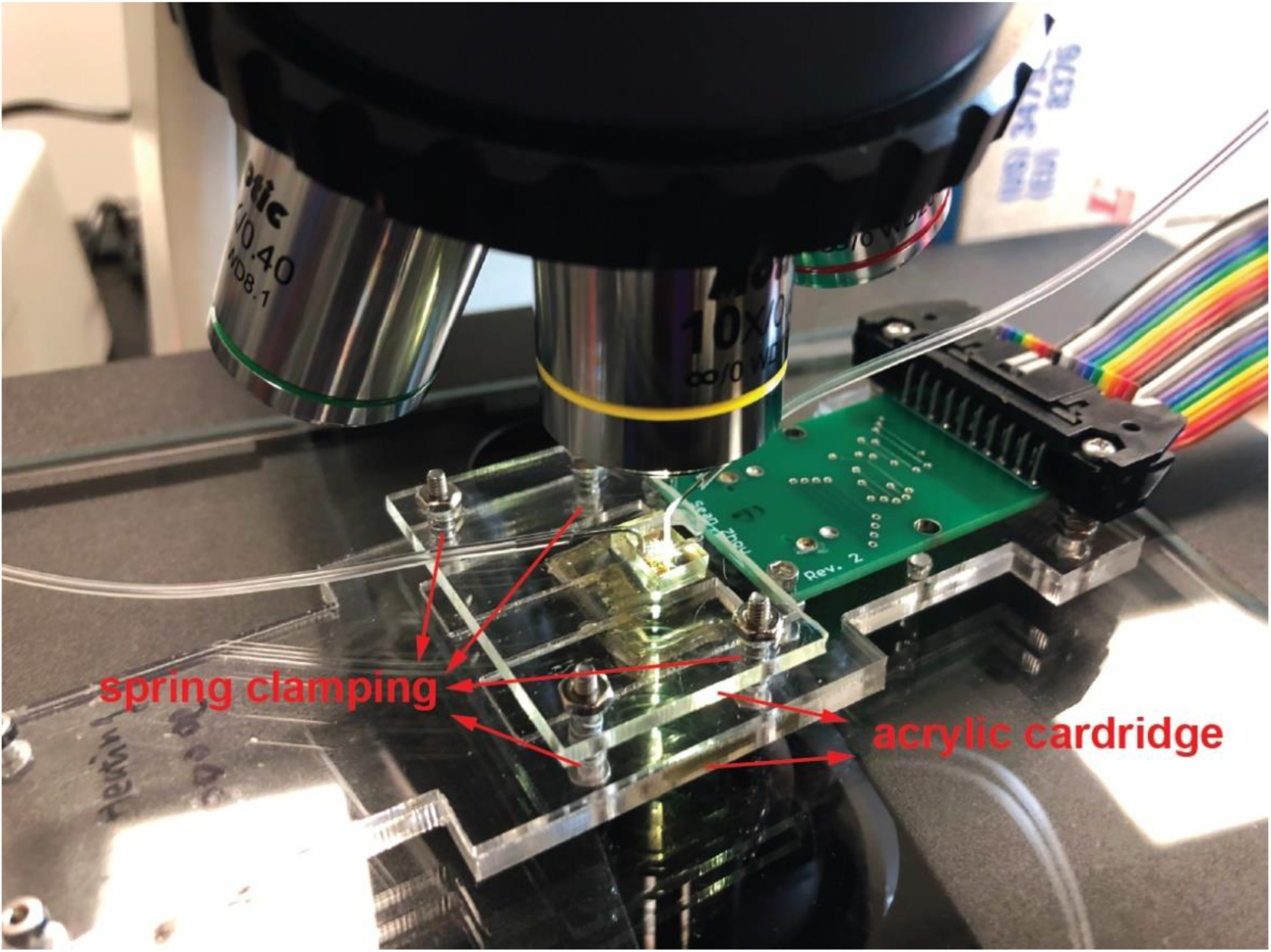
Holder showing spring-clamping. Acrylic plates with spring clamping enhance the sealing and prevent the system from leaking.

**Supplemental Figure 8.**
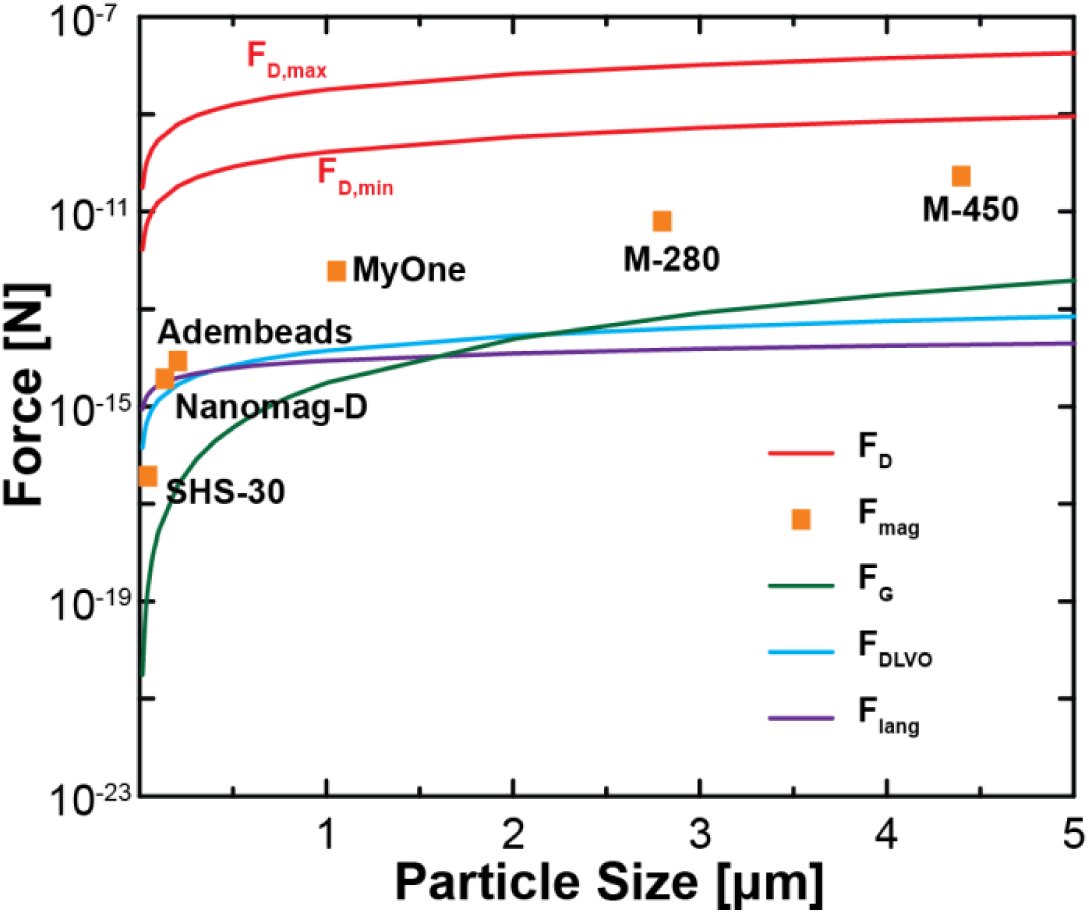
Hydrodynamic analysis. For sub-micron-sized MNPs (Adembeads, 200nm; Nanomag-D, 130nm; and SHS-30, 40nm), the magnetic force is comparable to DLVO forces and Langevin force. While Drag force was kept dominant over other forces in the setup to allow samples flowing at middle of the channel and thus extract the multi-parametric information, as shown in the figure, drag force is at least one decade larger than magnetic force even flowing the largest MNP (M-450, 4.5 μm).

**Supplemental Figure 9.**
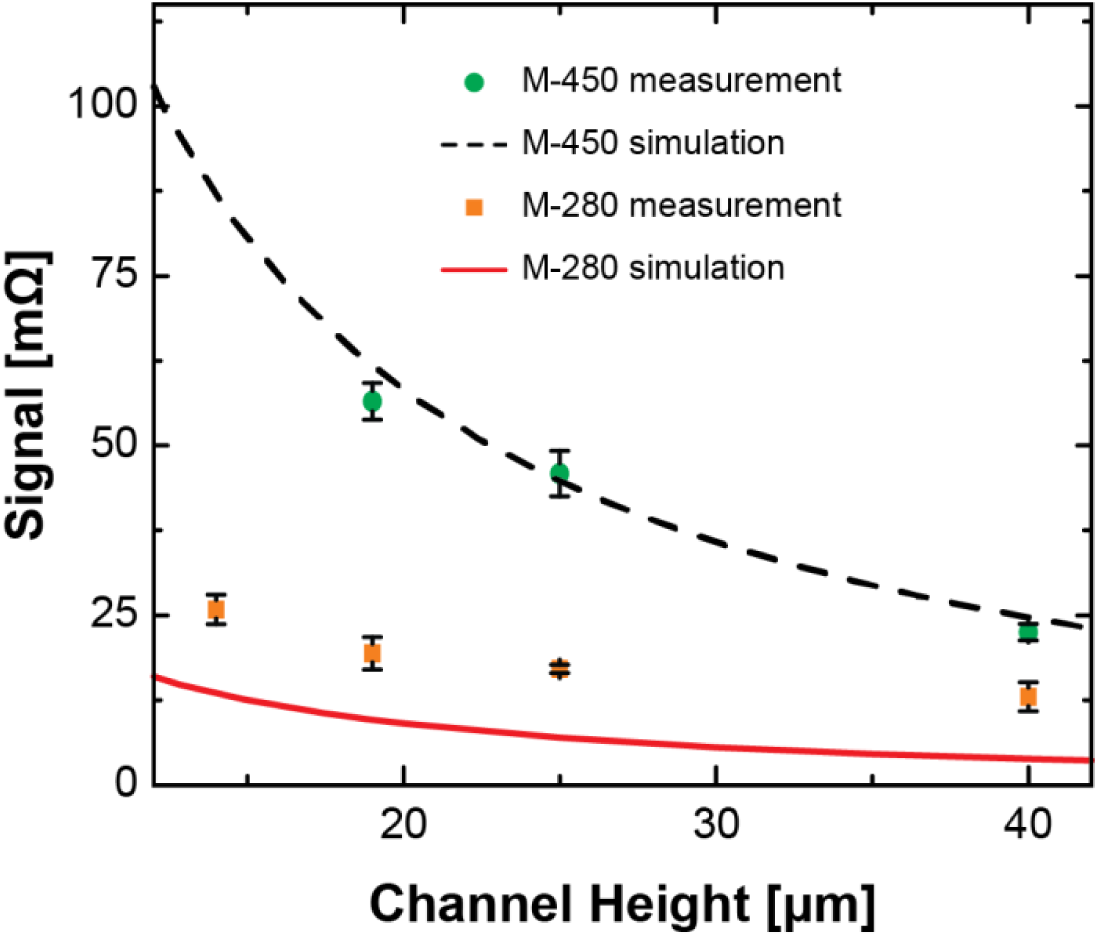
Comparison of measured results and simulations using Dynabeads (M-450 and M-280). The experimental result of M-450 is in excellent agreement with the simulation, while the average signal amplitude of M-280 deviated from the theoretical prediction owing to the aggregation and/or chaining.

**Supplemental Figure 10.**
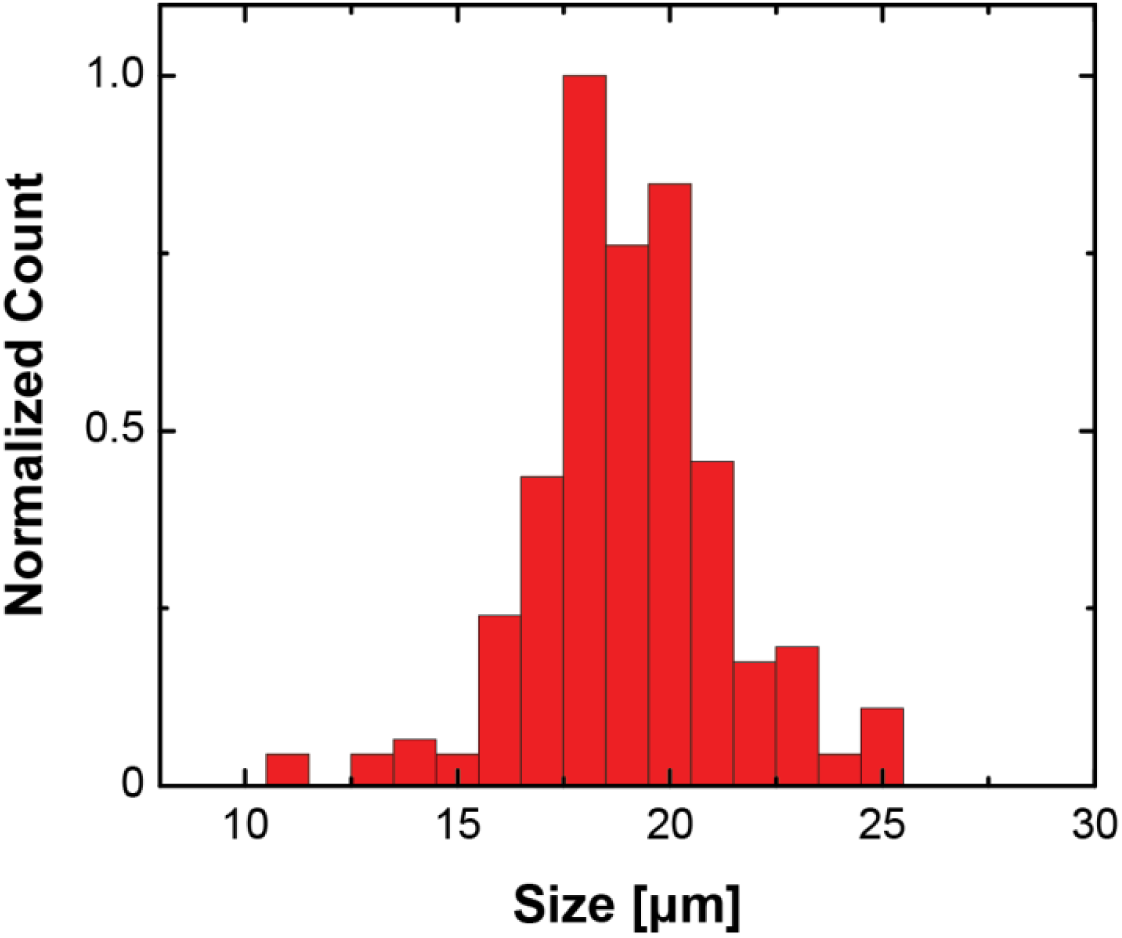
Size distribution of Panc-1 cancer cells. The cell size of Panc-1 cells was calculated using a Vi-CELL XR Cell Viability Analyzer (Beckman Coulter). The mean cell diameter size was calculated to be 19.51 µm.

**Supplemental Figure 11.**
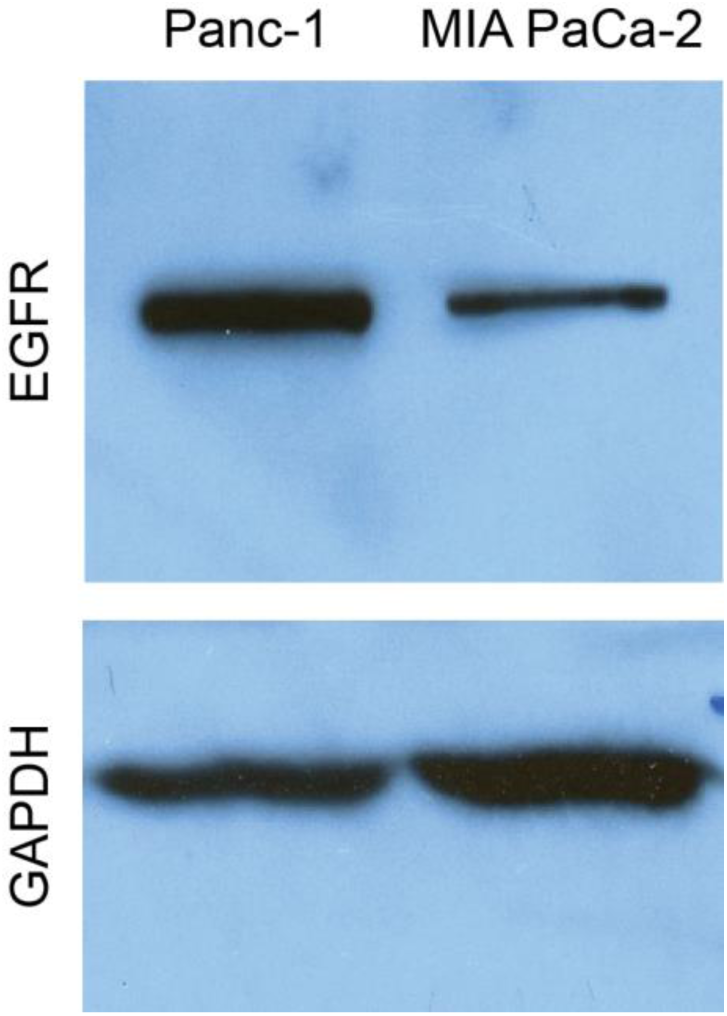
Panc-1 and MiaPaCa-2 cell lysates were subjected to Western blot analysis using anti-EGFR antibody. Glyceraldehyde 3-phosphate dehydrogenase (GAPDH), a house keeping protein, was used as the loading control. Panc-1 cells expressed more EGFR as compared to the MiaPaca-2 cells.

**Supplemental Figure 12.**
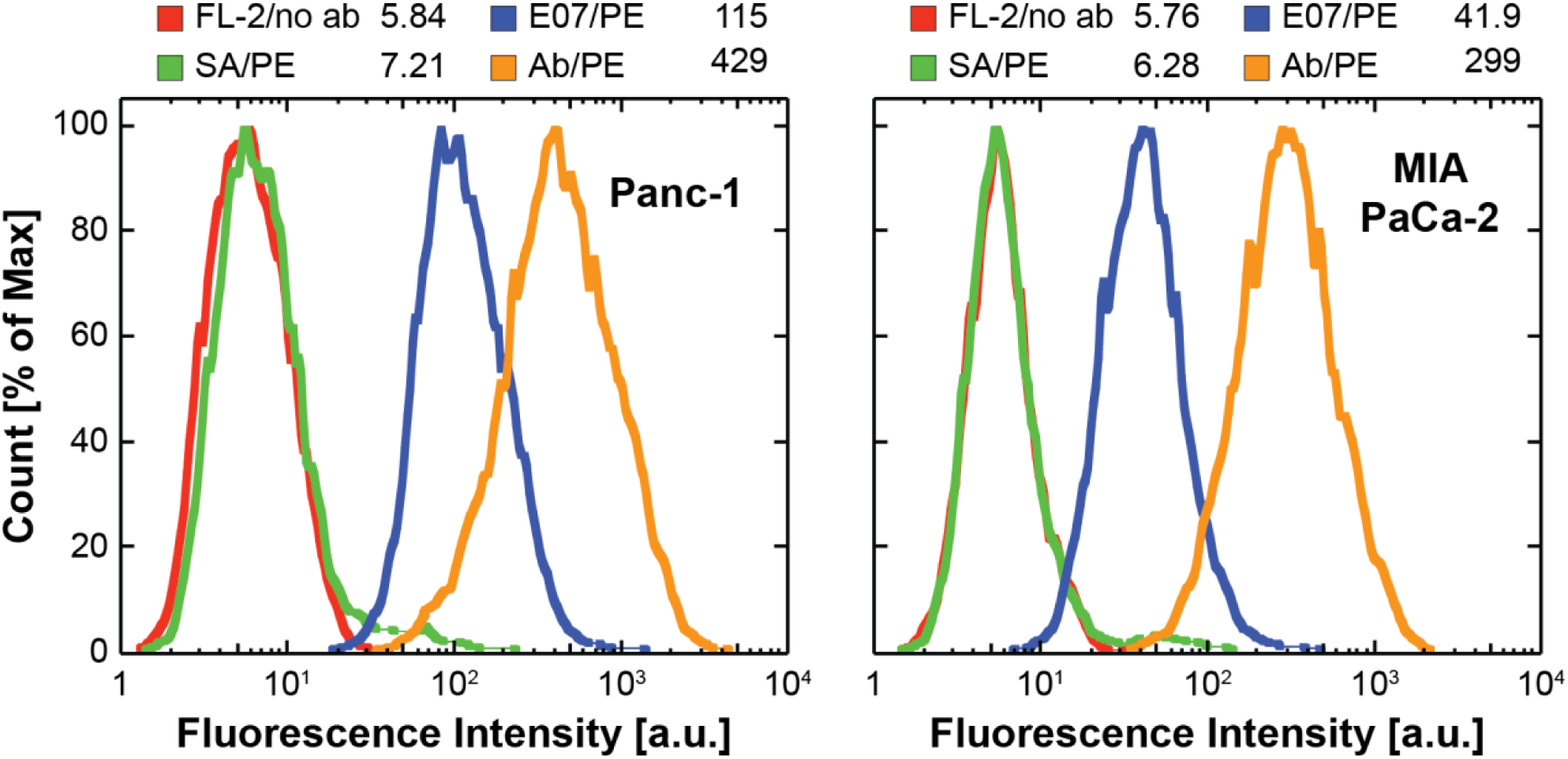
Optical FCM data for Panc-1 and MiaPaCa-2 with E07 aptamer and anti-EGFR antibody. Optical FCM analyses of anti-EGFR aptamer (E07) and antibody binding to the Panc-1 and MiaPaCa-2 cells were performed by treating the cells with the fluorophore phycoerythrin (PE)-conjugated 5’Biotin-E07 aptamer (blue) or PE-conjugated Biotin-anti-EGFR antibody (orange). Streptavidin-PE was used as the negative binding control (green). The data was analyzed, and the mean fluorescence intensity (MFI) was calculated from the histogram plots.

**Supplemental Figure 13.**
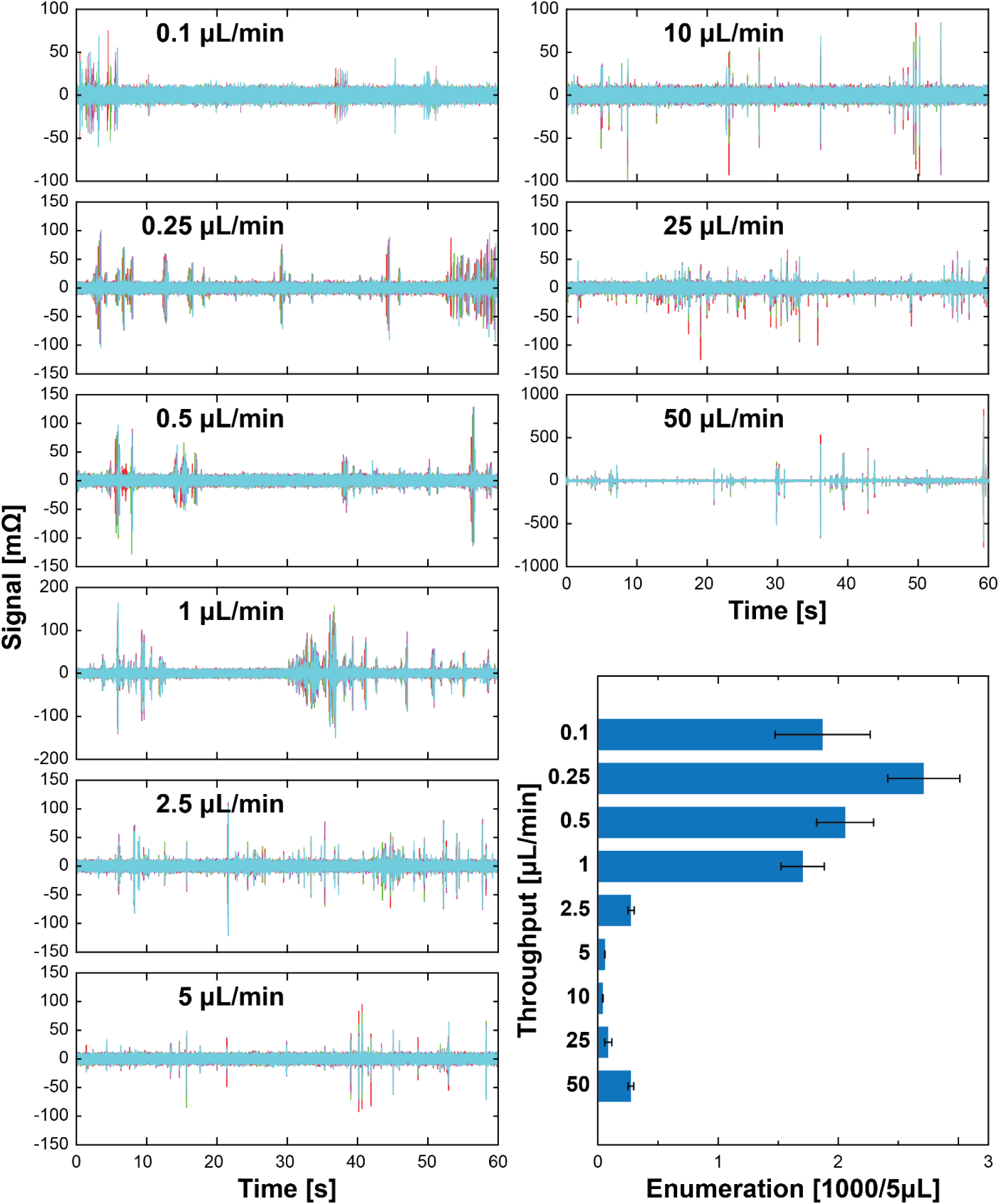
MFC measurements through varying throughputs across four sensors. The E07-decorated Panc-1 cells were measured in culture media using the same procedure. The data shows successful enumeration across two decades of throughput, ranging from 0.1 to 50 µL/min. Due to the viscosity (Momen-Heravi et al., 2012), the enumeration (*n* = 2711) and the best throughput (0.25 µL/min) for measurements were different from what was measured in **Fig. 5B** (*n* = 7140 under 0.1 µL/min). The distorted signal and sample aggregation occurred more frequently when throughput was increased, while Panc-1 cells were still also measurable (*n* = 273) under 50 µL/min of throughput.

**Supplemental Figure 14.**
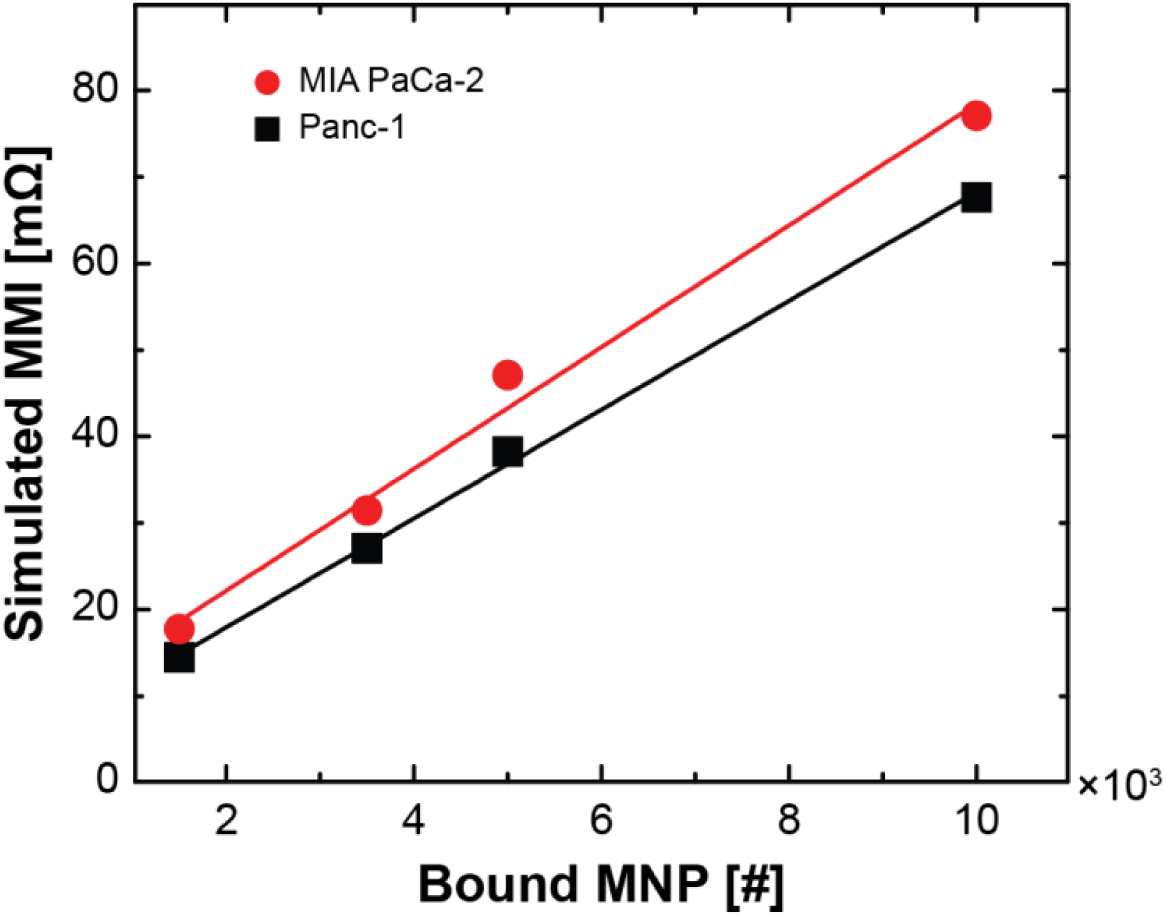
Simulated magnetic signal vs. number of bound MNPs to both Panc-1 and MiaPaCa-2 cells. In the simulation, the cell was regarded as a sphere with random distribution of surface EGFR, and each EGFR can be bound with one MNP. Due to the smaller size of MiaPaCa-2 (mean diameter = 16.71 μm) and proximity sensing of magnetism, its signal would be larger than Panc-1 if they have the same amount of bound MNPs.

